# Keratin isoform shifts modulate motility signals during wound healing

**DOI:** 10.1101/2023.05.04.538989

**Authors:** Benjamin A Nanes, Kushal Bhatt, Rajaa Boujemaa-Paterski, Evgenia Azarova, Sabahat Munawar, Divya Rajendran, Tadamoto Isogai, Kevin M Dean, Ohad Medalia, Gaudenz Danuser

## Abstract

Keratin intermediate filaments form strong mechanical scaffolds that confer structural stability to epithelial tissues, but the reason this function requires a protein family with 54 isoforms is not understood. During skin wound healing, a shift in keratin isoform expression alters the composition of keratin filaments. How this change modulates cellular function to support epidermal remodeling remains unclear. We report an unexpected effect of keratin isoform variation on kinase signal transduction. Increased expression of wound-associated keratin 6A, but not of steady-state keratin 5, potentiated keratinocyte migration and wound closure without compromising epidermal stability by activating myosin motors. This pathway depended on isoform-specific interaction between intrinsically disordered keratin head domains and non-filamentous vimentin shuttling myosin-activating kinases. These results substantially expand the functional repertoire of intermediate filaments from their canonical role as mechanical scaffolds to include roles as isoform-tuned signaling scaffolds that organize signal transduction cascades in space and time to influence epithelial cell state.

## Introduction

Effective tissue barrier formation depends on keratin intermediate filaments^1^. Joined through desmosomes, keratin filaments form strong trans-cellular networks, which function as mechanical scaffolds^2^. Fifty-four different keratin isoforms are expressed in an intricate pattern depending on tissue, anatomic site, and differentiation state^3^. Keratin filaments are particularly important in the skin, where their disruption causes epidermal fragility in a number of diseases^4^, and specific keratin isoform mixtures mark epidermal layers and associated cell states (Figure 1A) ^1^. However, epidermal stability must be balanced with plasticity^1^. Too much stability relative to plasticity can limit remodeling, resulting, for example, in non-healing wounds^5^. Too much plasticity relative to stability can result in uncontrolled or abnormal remodeling, a feature of numerous skin diseases ranging from cancer to psoriasis^6,7^. The molecular mechanisms underlying the balance between epidermal stability and plasticity largely remain to be uncovered.

**Figure 1.**
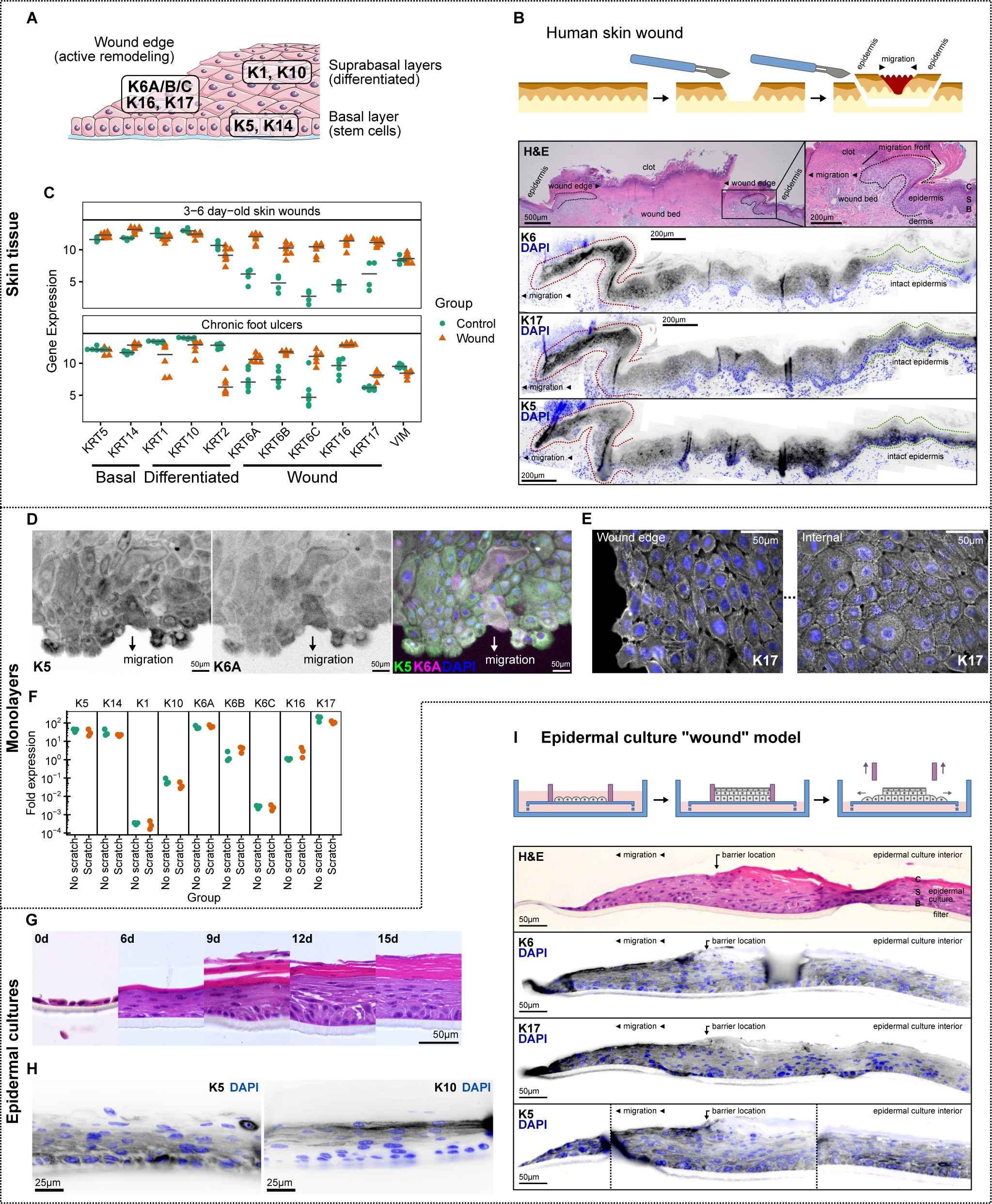
Epidermal remodeling triggers a keratin isoform switch. A. Diagram of keratin isoforms expressed in steady-state and actively remodeling epidermis. B. Top: Schematic diagram of a skin excision following a diagnostic biopsy, which includes healing edges from the biopsy wound. Middle: Hematoxylin and eosin (H&E)-staining. Dotted outlines, epidermal migration fronts. Epidermal layers: B, basal; S, spinous (suprabasal); C, cornified. Bottom: Immunofluorescence labeling of wound-associated K6 and K17, and steady-state basal epidermal K5. Red dotted outline, epidermal migration front at the wound edge; Green dotted outline, intact epidermis away from the wound. C. Intermediate filament expression in acute (Top, GSE97615) and chronic (Bottom, GSE80178) human skin wounds and intact skin controls. D-E. Confluent keratinocyte monolayers were scratched with a pipette tip to create wounds, allowed to migrate for 24-hours, then fixed, labeled for immunofluorescence of steady-state K5 and wound-associated K6 (D) or K17 (E), and imaged. In (E), images of the migration edge and monolayer interior were acquired with the same illumination and detection settings. I. F. Keratinocyte monolayers were preserved intact or repeatedly scratched with a pipette tip (see Methods), then processed for RNA isolation and quantification of keratin transcript levels by qRT-PCR. Individual keratin transcript levels were compared to the average level of all keratin transcripts measured. *n* = 3 monolayers per group. G-H. Keratinocytes were cultured at an air-liquid interface to induce stratification. G, Hematoxylin and eosin (H&E) staining after different growth periods. H, Immunofluorescence labeling of a basal epidermal keratin, K5, and a differentiated epidermal keratin, K10. I. Top: Schematic diagram of the epidermal culture wound model. Removal of a barrier mold allows keratinocytes from the stratified culture to migrate into the “wound” area (see Methods). Middle: H&E staining after 72 hours of migration. Arrow, prior location of the barrier. Epidermal layers: B, basal; S, spinous (suprabasal); C, cornified. Bottom: Immunofluorescence labeling of wound-associated K6 and K17, and a steady-state basal K5. Left, migration edge; Right, epidermal culture interior. Dashed vertical lines, image re-alignment correcting for folds in the tissue section. See also Figure S1B.

During wound healing, the required balance between epidermal stability and plasticity shifts, as keratinocytes change shape, migrate into the wound area, and reform the epidermal architecture^1^. These tissue-level processes are accompanied at the cellular scale by increased expression of five *wound-associated keratin* isoforms: keratin 6A (K6A), K6B, K6C, K16, and K17 (Figure 1A) ^8–10^. Expression of wound-associated keratins also increases in other active remodeling epidermal states, including psoriasis^11,12^, squamous cell carcinoma^13^, hypertrophic scars^14^, and following tissue expander placement^15^.

While expression of wound-associated keratin isoforms is a reliable marker of an active remodeling state, determining whether these keratin isoforms have specific cellular functions that support wound healing has been experimentally and conceptually challenging. Depleting or mutating individual keratins frequently disrupts keratin filament architecture and the more general mechanical scaffold function^16^. Partial redundancy between keratin isoforms further complicates the interpretation of monogenic models^17^. Wound-associated keratin knock-out mice often demonstrate prominent defects in epidermal integrity and keratinocyte cell-cell adhesion, but lack generalized wound-healing defects^18–21^. Similarly, disease-causing mutations in human wound-associated keratin genes broadly disrupt epidermal stability where the affected isoform is expressed^22,23^. Thus, while wound-associated keratins contribute to the canonical intermediate filament mechanical scaffold, it is less clear if additional functions of these isoforms specifically promote wound closure.

In order to disentangle the specific contributions of wound-associated keratins to epidermal remodeling from the general function of keratin mechanical scaffolds, we took a gain-of-function rather than a loss-of-function approach. We found that increased expression of wound-associated K6A, but not steady-state K5, potentiated keratinocyte migration without compromising cell-cell adhesion. Rather than altering the mechanical response to external force, K6A expression increased cellular force generation via activation of actomyosin contractility. This role in signal transduction was mediated by preferential interaction of the K6A intrinsically disordered head domain with non-filamentous vimentin. K6A-enriched keratin filaments and non-filamentous vimentin transiently anchored a signaling complex, spatially organizing regulatory kinases and myosin subunits to promote myosin activation. Thus, by wound-induced keratin isoform switching, epidermal cells amplify and spatially organize a signal transduction cascade that promotes critical functions for the wound healing process. These results highlight the complex interplay between the cytoskeleton and apparently diffuse pools of cytoplasmic proteins to spatiotemporally structure biochemical reactions.

## Results

### An epidermal culture model of wound-induced keratin switching

Keratin expression change following wounding has been observed in animal models and humans^8–10^. We reproduced these behaviors in a human skin excision specimen, where we detected increased expression of wound-associated K6 and K17 at healing wound edges (Figure 1B). We also reanalyzed published transcriptomics data of acute^24^ and chronic wounds^25^ with a focus on intermediate filament expression and found strongly increased expression of all five wound-associated keratin genes (Figure 1C). However, wound-induced keratin switching has not previously been observed in cultured keratinocytes. Furthermore, cultured keratinocytes often express wound-associated keratins at baseline^26^, possibly reflecting the increased proliferative rate of keratinocytes in culture^27^. This has complicated efforts to understand the implications of keratin switching at the cellular and subcellular levels.

Consistent with prior reports^28,29^, we found that an hTERT and Cdk-4 immortalized human keratinocyte cell line grown in submerged monolayer culture expressed steady-state basal level keratins K5 and K14 and wound-associated K6 and K17 (Figure S1A). Steady-state differentiated keratins K1 and K10 were barely detectable (Figure S1A), as expected for a basal keratinocyte phenotype. Immunofluorescence of monolayer scratch wounds did not show any change in expression of K6 or K17 relative to K5 at the wound edge (Figure 1D-E). Similarly, bulk qRT-PCR did not show any change in wound-associated keratin transcript levels in scratched keratinocyte monolayers compared to unwounded controls (Figure 1F).

While keratin expression remained stable in submerged monolayer culture, keratinocytes at an air-liquid interface formed a multilevel structure undergoing differentiation, with increased expression of K10 in the upper layers (Figure 1G-H), closely resembling the stratified squamous architecture of the epidermis^30^. We leveraged these stratified epidermal cultures to create a three-dimensional wound model. Cultures were grown within a barrier mold, followed by barrier removal to allow keratinocyte migration (Figure 1I). In stark contrast to monolayer scratch wounds, wound-associated K6 and K17 expression was notably increased at the migrating edge of stratified cultures (Figures 1I and S1B). The presence of wound-induced keratin switching in the epidermal culture model prompted us to more closely examine the function of wound-associated keratins in this system.

### Wound-associated K6A supports epidermal migration

We sought an interventional strategy for modulating relative levels of wound-associated versus steady-state keratins. Prior attempts to knock down or delete keratin genes have resulted in prominent loss of cell-cell adhesion and decreased mechanical integrity^31–35^, a finding we reproduced with even heterozygous deletion of KRT6 isoforms (Figure S1C-D). To circumvent such global effects, we took a gain-of-function approach and co-expressed fluorescently tagged K6A, a wound-associated keratin, and K5, the closest steady-state paralog to K6A with 80% amino acid sequence identity, and used fluorescence activated cell sorting to isolate cell populations with different relative keratin expression (Figure 2A).

**Figure 2.**
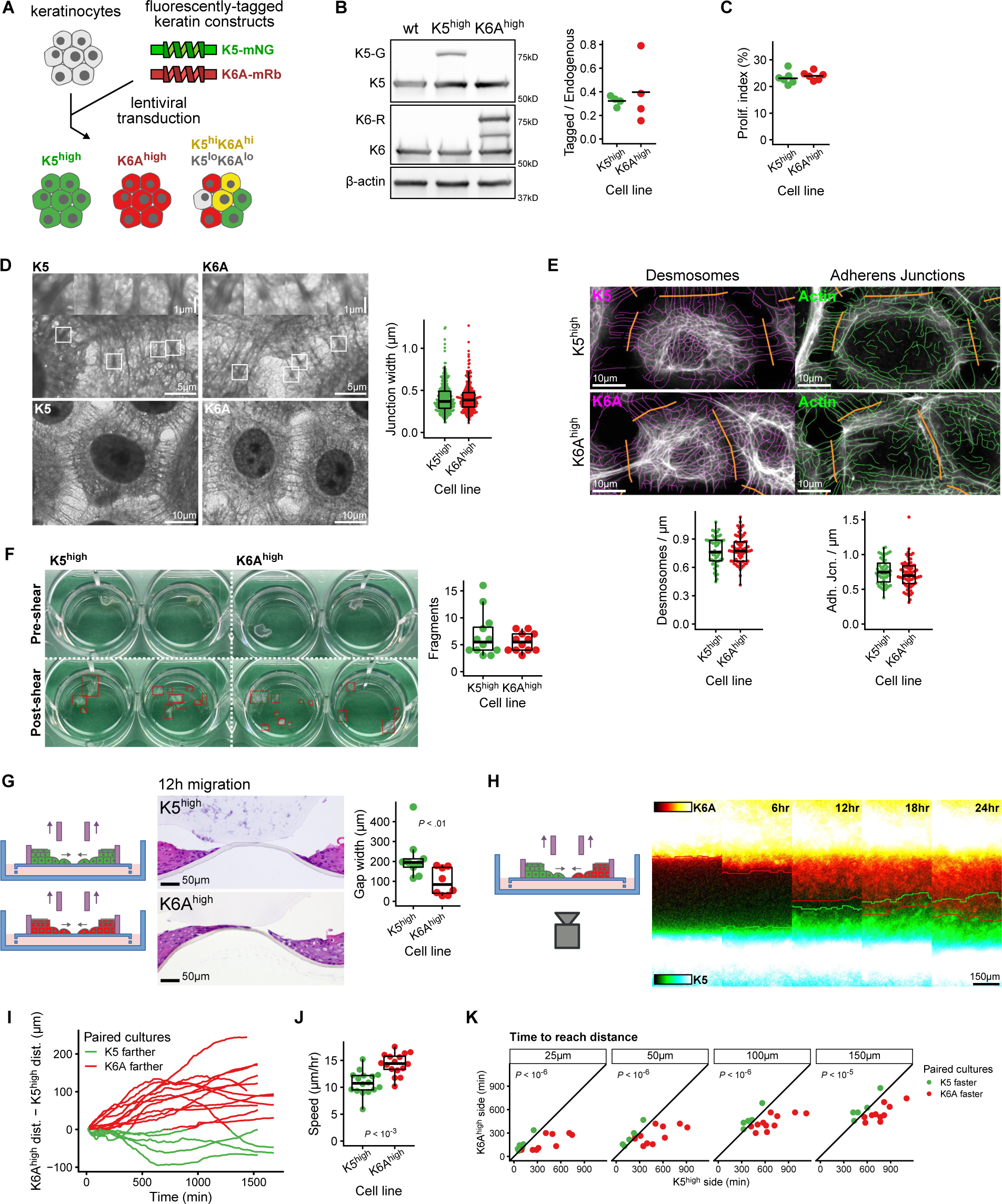
Wound-associated K6A supports epidermal migration without compromising mechanical stability. A. Schematic diagram of the gain-of-function keratin expression model. Cultured keratinocytes were transduced with lentivirus to express tagged K5 and K6A, then divided by fluorescence activated cell sorting into different populations including those with high levels of K5 (K5^high^), high levels of K6A (K6A^high^), high levels of both keratins (K5^hi^K6A^hi^), and low levels of both keratins (K5^lo^K6A^lo^). B. Western blot analysis of endogenous (K5, K6) and exogenously expressed (K5-G, K6-R) keratins in parental (wt), K5^high^, and K6A^high^ cell lines (see also Figure S1E). Right: Ratio of K5-G or K6-R to K5 or K6. *n* = 4 samples per group. C. Proliferation of K5^high^ and K6A^high^ cells in culture measured by uptake of 5-ethynyl-2’-deoxyuridine (EdU) during a 60-minute incubation. Proliferation index, defined as the percentage of nuclei with EdU incorporation. *n* = 6 samples per group. D. Transmission electron micrographs of K5^high^ and K6A^high^ monolayers. Insets highlight typical electron-dense cell-cell junction complexes. Right, distribution of junction complex widths. *n* = 314-348 junctions from 40-41 images per group. E. Fluorescence imaging of K5^high^ and K6A^high^ cell monolayers with partial segmentation of keratin and actin filament networks. Desmosomes and adherens junctions were identified, respectively, by keratin and actin filament bridges between adjacent cells. *n* = 55-71 cell-cell borders from 17-18 images per group. F. Mechanical stability of keratinocyte monolayers measured using a dispase-based fragmentation assay (see Methods). *n* = 12 monolayers per group. G. Paired epidermal cultures separated by a 500-μm barrier were grown from K5^high^ and K6A^high^ cells. Following removal of the barrier and 12-hours of migration, the remaining gap width was measured in H&E-stained tissue sections. *n* = 8-9 epidermal culture pairs per group. H-K. Live imaging of epidermal migration. Paired epidermal cultures separated by a 500-μm barrier were grown with K5^high^ and K6A^high^ cell lines on opposite sides of each barrier. Following barrier removal, cells were imaged over 24 hours (H; see also Video 1; see Methods). Red and green lines, farthest migration extent of the K6A^high^ and K5^high^ culture halves at each time point. I, Difference in migration distance, defined as farthest distance migrated by the K6A^high^ half minus farthest distance migrated by the K5^high^ half. Each line represents one culture pair. J, Average migration speed over 20-hours, defined as farthest migration distance divided by total time. K, Time needed to reach the designated distance checkpoint. Dots represent culture pairs, with x- and y-axis location corresponding to time required for the K5^high^ and K6A^high^ halves to reach the designated distance. Dots blow the diagonal represent culture pairs where the K6A^high^ half reached the designated distance checkpoint first; dots above the diagonal represent culture pairs where the K5^high^ half reached the checkpoint first. *n* = 16 epidermal culture pairs.

We created K5^high^/K6A^low^ (K5^high^) and K5^low^/K6A^high^ (K6A^high^) cell lines with moderately increased (approximately 35%) levels of the keratin marked “high” (Figures 2B and S1E), reasoning that a small change in keratin expression could avoid filament network disruption observed with higher levels of overexpression. ^36,37^ Because cells maintain an equimolar balance between type-I and type-II keratin isoforms, which polymerize as obligate heterodimers, we expect that modestly increasing K5 or K6A protein levels, both type-II keratins, may have resulted in small compensatory shifts of endogenous keratins, distributed over all expressed isoforms. ^3^ However, we confirmed that endogenous keratin protein and transcript levels were similar in K5^high^ and K6A^high^ cells (Figure S1F-G), indicating that comparison of these cell lines reflects directly engineered keratin expression changes, rather than indirect compensation. Importantly, increasing K5 or K6A protein levels did not affect cell proliferation (Figure 2C), cell junction morphology (Figures 2D and S1H), cell junction number (Figure 2E), or mechanical integrity as measured by monolayer fragmentation assays (Figure 2F).

We then proceeded to test the migration potential of K5^high^ and K6A^high^ keratinocytes. In three-dimensional epidermal migration assays, K6A^high^ epidermal cultures migrated farther than K5^high^ epidermal cultures over a 12-hour period (Figure 2G). We further took advantage of the bright fluorescence signals from the tagged keratins to enable live imaging of paired K5^high^ and K6A^high^ epidermal cultures following removal of a separating barrier (Figure 2H). While the partially translucent polycarbonate filter limited image quality, the fluorescence signal was sufficient to identify and track the migration edges using custom software (Figure 2H and Video 1) ^38^. We observed that K6A^high^ epidermal cultures moved faster into the gap than their K5^high^ counterparts, and K6A^high^ cultures maintained their migration advantage over a 20-hour migration period (Figure 2I-K). Therefore, increased expression of wound-associated K6A offers a significant benefit to wound closure in three-dimensional epidermal cultures.

### Wound-associated K6A supports migration in monolayers and individual cells

In order to perform high-resolution imaging of potential differences in cytoskeletal organization and dynamics between K5^high^ and K6A^high^ cells, we turned to a monolayer wound healing assay. In contrast to the three-dimensional epidermal wound model, we were initially unable to identify a clear difference in migration speed between K5^high^ and K6A^high^ monolayers (Figure S2A-D and Video 2). Since the monolayer migration assay had significant wound-to-wound variability (Figure S2B), we wondered whether subtle but systematic differences in migration behavior between individual K6A^high^ and K5^high^ cells were missed in the coarser population level analysis.

To test this possibility, we created a mosaic population of keratinocytes including K5^high^, K6A^high^, K5^high^/K6A^high^, and K5^low^/K6A^low^ cells and performed live imaging of the monolayer wound healing response (Figure 3A and Video 3). We applied particle image velocimetry to track local migration throughout the monolayer (Figure 3B, bottom; Video 3, right) ^38^ and segmented the monolayers into different keratin expression regions (Figure 3B, top; Video 3, left), permitting comparison of migration speeds between keratin expression regions within the same wound and controlling for between-wound variability.

**Figure 3.**
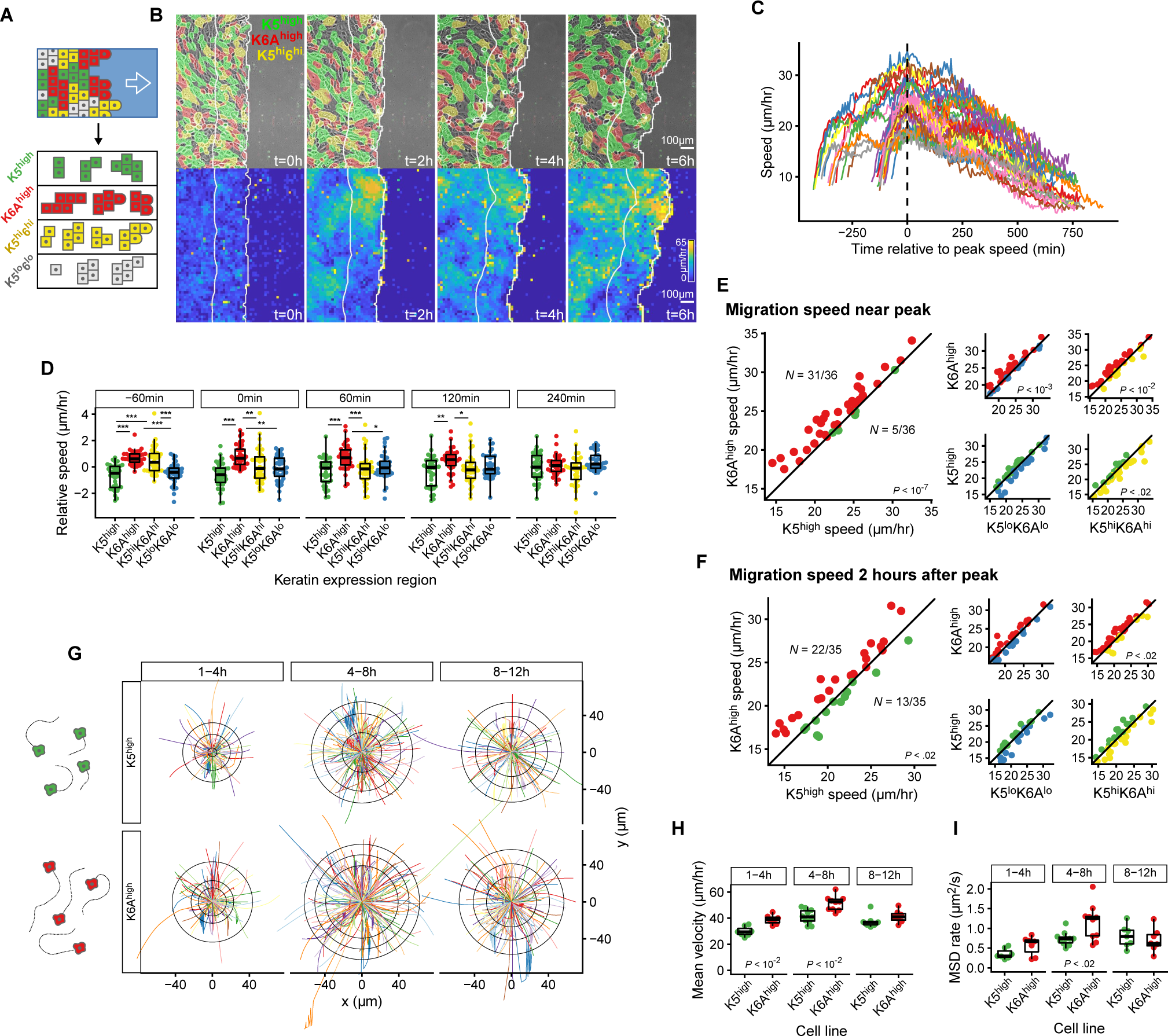
Wound-associated K6A supports keratinocyte migration in monolayers and single cells. A. Schematic diagram of a mosaic monolayer containing K5^high^ (green), K6A^high^ (red), K5^high^/K6A^high^ (K5^hi^6^hi^, yellow), and K5^low^/K6A^low^ (K5^lo^6^lo^, grey) cells during migration. Local migration speeds are compared between different keratin expression regions. B. Top: Keratin expression regions. Bottom: Local migration speeds calculated using a computer vision pipeline (see Methods). White lines, 200-μm band tracking the wound edge. See also Video 3. C. Time courses of average local migration speed 10-μm to 200-μm from the wound edge aligned relative to the global peak in migration speed. *n* = 36 scratch wounds. D. Relative local migration speed within each keratin expression region at different time points relative to the peak in migration speed. Relative local speeds are defined as the local migration speed for a keratin expression region minus the average migration speed across all regions 10-μm to 200-μm from the migration edge. *n* = 36 scratch wounds. E-F. Pairwise comparison of average local migration speed between keratin expression regions at the peak in migration speed (E) and 2-hours later (F). Each point represents one scratch wound, with x- and y-axis location corresponding to the migration speed in different keratin expression regions. *n* = 36 scratch wounds. See also Video 4. G-I. Movement of sparsely seeded, individual K5^high^ and K6A^high^ keratinocytes. G, Cell trajectories aligned at the initial position, including the first hour of each track within the time window. Rings, quantiles of maximum distance from the origin. See also Video 5. H-I, Mean track velocity (H) and mean squared displacement rate (I) in K5^high^ and K6A^high^ cells. *n* = 7-10 movies with 144-232 tracks per group.

All wounds followed a similar pattern. After scratching the monolayer with a pipette tip, the overall migration speed rapidly increased to a peak followed by slow deceleration, though the height and timing of the peak varied between experimental repeats (Figure 3C). Around the time of the overall migration speed peak, K6A^high^ regions exhibited a small, but consistent, migration advantage (Figure 3D). In the large majority (31 of 36) of wounds, K6A^high^ regions had higher speeds than K5^high^ regions (Figure 3E). K6A^high^ regions also had higher speeds than K5^high^/K6A^high^ and K5^low^/K6A^low^ regions, and K5^high^/K6A^high^ regions had higher speeds than K5^high^ regions, indicating a dose-dependent effect (Figure 3E). Interestingly, the K6A^high^ migration advantage was transient. By 2 hours after the migration peak, K5^high^ region speeds began to approach those of the K6A^high^ regions (Figure 3F and Video 4). Consistent with these results, K6A^high^ cells were modestly enriched near the wound edge of a mosaic population monolayer fixed after 18-hours of migration (Figure S2E).

Given that K5^high^ and K6A^high^ keratinocytes retain similar cell-cell adhesion properties (Figure 2D-F), we asked whether the K6A^high^ migration advantage was solely an emergent property of collective migration or if it could also be detected in individual keratinocytes. To address this question, we seeded K5^high^ and K6A^high^ cells at low density and tracked their movement over 12-hours using live imaging (Figure 3G and Video 5). Much like in the monolayer migration assays, K6A^high^ cells displayed a small advantage in migration speed and displacement, which peaked 4 to 8-hours after seeding (Figure 3G-I). Also similar to monolayer migration, the migration advantage was temporary, with the differences between K5^high^ and K6A^high^ cells becoming less apparent by 8 to 12-hours after seeding (Figure 3G-I). Thus, wound-associated K6A transiently potentiates keratinocyte migration at least in part through a cell autonomous effect.

### Wound-associated K6A alters keratin filament dynamics

To address how the relative levels of keratin isoforms affect migration, we examined possible differences in keratin filament organization between K5^high^ and K6A^high^ cells. We imaged keratin filaments over time and used a computer vision pipeline to segment the filament networks (Figures 4A-B, S3A, and Video 6)^39^. In snapshots, the K5^high^ and K6A^high^ filament networks were similar. K5^high^ filament networks appeared to contain a somewhat higher number of very short filaments (Figure S3B), and an associated slight increase in overall filament density (Figure S3C), compared to K6A^high^ networks, possibly reflecting a slight difference in the imaging properties of the two fluorescent tags. Filament curvature was indistinguishable between K5^high^ and K6A^high^ cells (Figure S3D-F). However, when analyzed over time, K5^high^ filaments displayed larger intracellular displacements than K6A^high^ filaments (Figure 4A-B).

**Figure 4.**
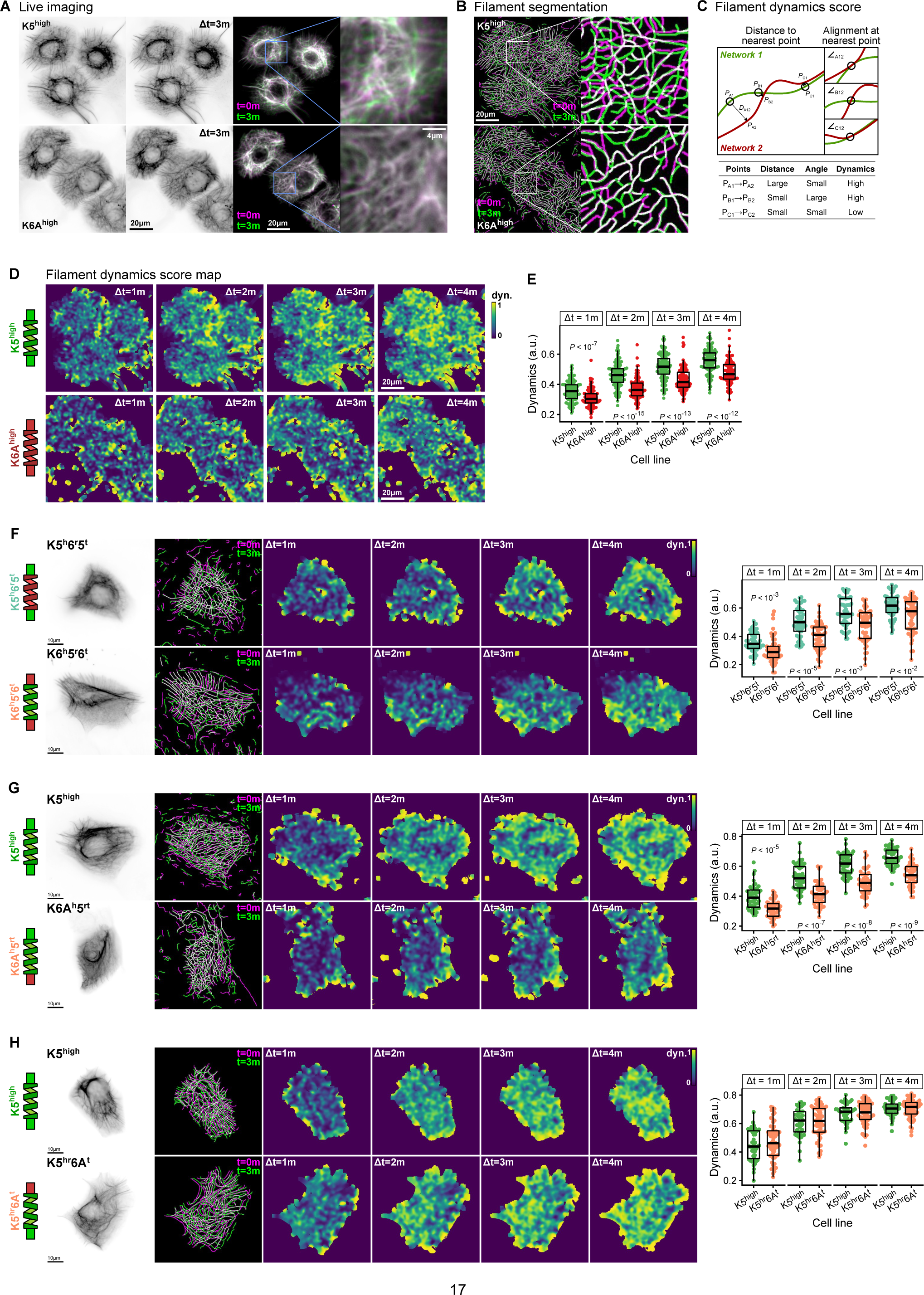
Wound-associated K6A alters keratin filament network dynamics. A. Keratin filaments in K5^high^ and K6A^high^ cells. Right, pseudo-color overlay of filament images captured 3 minutes apart. B. Filament networks segmented using a computer vision pipeline (see Methods). As in (A), overlay of networks 3 minutes apart displays the dynamics of the network architecture. See also Video 6. C. Schematic diagram of filament dynamics score calculation (see Methods). D. Smoothed maps of local filament dynamics scores over different time intervals in K5^high^ and K6A^high^ cells. E. Spatially averaged filament dynamics scores for each cell. *n* = 110-139 cells in 24-25 movies per group. F. Comparison of filament dynamics scores between keratinocyte cell lines expressing chimeric keratins containing the head and tail domains of K5 or K6A joined to the rod domain of the opposite keratin (K5^h^6^r^5^t^ and K6^h^5^r^6^t^). *n* = 51-52 cells in 21-29 movies per group. G-H. Comparison of filament dynamics scores between keratinocyte cell lines expressing chimeric keratins containing only the head (G; K6A^h^5^rt^) or tail (H; K5^hr^6A^t^) domains of K6A joined to the remainder of K5 with K5^high^ cells. G, *n* = 44-47 cells in 28 movies per group. H, *n* = 45-60 cells in 37-39 movies per group.

To quantify differences in filament network dynamics, we created a metric reflecting changes in position and orientation of filaments between timepoints at one-minute intervals^39^. Each point on the filament network at baseline is scored by the distance and difference in orientation of the nearest filament in the next considered timepoint (Figure 4C). We then spatially smoothed the local scores to generate continuous filament dynamics score maps and averaged over individual cells (Figure 4D). Comparing K5^high^ to K6A^high^ cells, we found that K6A systematically decreased intracellular dynamics of keratin filaments (Figure 4E). We confirmed that the difference in filament dynamics between K5^high^ and K6A^high^ cells could not be explained by the small differences in number of very short filaments or filament density (Figure S3G-H).

We further extended this approach using two-color live imaging of K5^high^, K6A^high^, and K5^high^/K6A^high^ cells, allowing us to segment and calculate filament dynamics scores for K5 and K6A simultaneously (Figure S3I-O and Video 6). We found a dose-dependent effect of K6A on filament dynamics, with the highest dynamics scores for K5^high^ cells, the lowest dynamics for K6A^high^ cells, and intermediate dynamics for K5^high^/K6A^high^ cells (Figure S3M-N). Furthermore, we could compare filament segmentations and dynamics scores between the K5 and K6A channels in K5^high^/K6A^high^ cells. Given the known propensity for individual keratins to intermix within filament networks^40^, K5 and K6A network segmentations overlapped considerably, as expected (Figure S3L and Video 6). Additionally, K5 and K6A dynamics scores for K5^high^/K6A^high^ cells were highly correlated, without evidence of bias between detection channels (Figure S3O), confirming the robustness of the filament segmentation algorithm and network dynamics score. Taken together, these results establish a direct relationship between the mixture of keratin isoforms assembled into filaments and the dynamics of the networks that those filaments form.

The filament dynamics score is a composite measure of both intrinsic properties of the filaments, such as stiffness, and extrinsic interactions between the filament network and other cellular components, such as the contractile actomyosin cytoskeleton. We reasoned that if a difference in contractility was responsible for the difference between K5^high^ and K6A^high^ filament dynamics, inhibiting myosin motors would reduce that effect. Indeed, we found that the specific myosin II inhibitor Blebbistatin completely eliminated the difference between K5^high^ and K6A^high^ filament dynamics (Figure S4A), while two upstream ROCK inhibitors had smaller effects (Figure S4B). While insufficient on their own to define a mechanism, these results establish a link between keratin filament dynamics and the actomyosin system.

We reasoned that determining the domain of the keratin protein responsible for the dynamics difference between K5^high^ and K6A^high^ filaments might further untangle this effect. Keratin proteins are composed of a central alpha helical rod domain flanked by intrinsically disordered head and tail domains^41,42^. We created chimeric keratin constructs composed of the head and tail domains of K5 or K6A fused to the rod domain from the opposite keratin (K5^h^6^r^5^t^ and K6^h^5^r^6^t^). Upon expressing each of these chimeras in cultured keratinocytes, we found that K6^h^5^r^6^t^ decreased filament dynamics compared to K5^h^6^r^5^t^ (Figure 4F). We further created chimeric keratins containing only the head or tail domain of K6A fused to the remainder of K5 (K6A^h^5^rt^ and K5^hr^6A^t^) and determined that the K6A head domain alone was sufficient to decrease filament dynamics (Figure 4G), while the K6A tail domain had no effect (Figure 4H). Thus, the intrinsically disordered K6A head domain, rather than the structural rod domain, is responsible for decreasing keratin filament dynamics. While we cannot formally exclude contributions of the head domain to the mechanical properties of filaments, this result, combined with the similarity of K5^high^ and K6A^high^ filament networks in snapshots (Figure S3B-F), suggests that the changes in filament dynamics between K5^high^ and K6A^high^ cells are more likely related to changes in cytoskeletal force generation than changes in filament stiffness.

### Wound-associated K6A increases cellular force generation by activating myosin

To directly test whether changing keratin filament composition affects cellular force generation, we compared the ability of K5^high^ and K6A^high^ cells to deform an elastic substrate using traction force microscopy^43^. We found that K6A^high^ cells generated increased traction forces and strain energy compared to K5^high^ cells (Figures 5A and S4C). Using alternate cell lines expressing short-tagged keratins, we confirmed that fluorescent tags were not responsible for this effect (Figure S4D). Furthermore, expressing the K6A^h^5^rt^ chimera also increased force generation (Figures 5B and S4E), indicating that, as with filament dynamics, traction force generation was modulated by differences in keratin head domains. Importantly, the increased strain energy, which measures the total work of contraction executed by the cell, cannot be explained by changes in material properties such as filament elasticity or differences in organization of cell-matrix adhesions. The latter would in fact be expected as a downstream effect secondary to increased traction stress. ^44,45^ This finding directly links shifts in the abundance of wound-associated K6A to cellular force generation.

**Figure 5.**
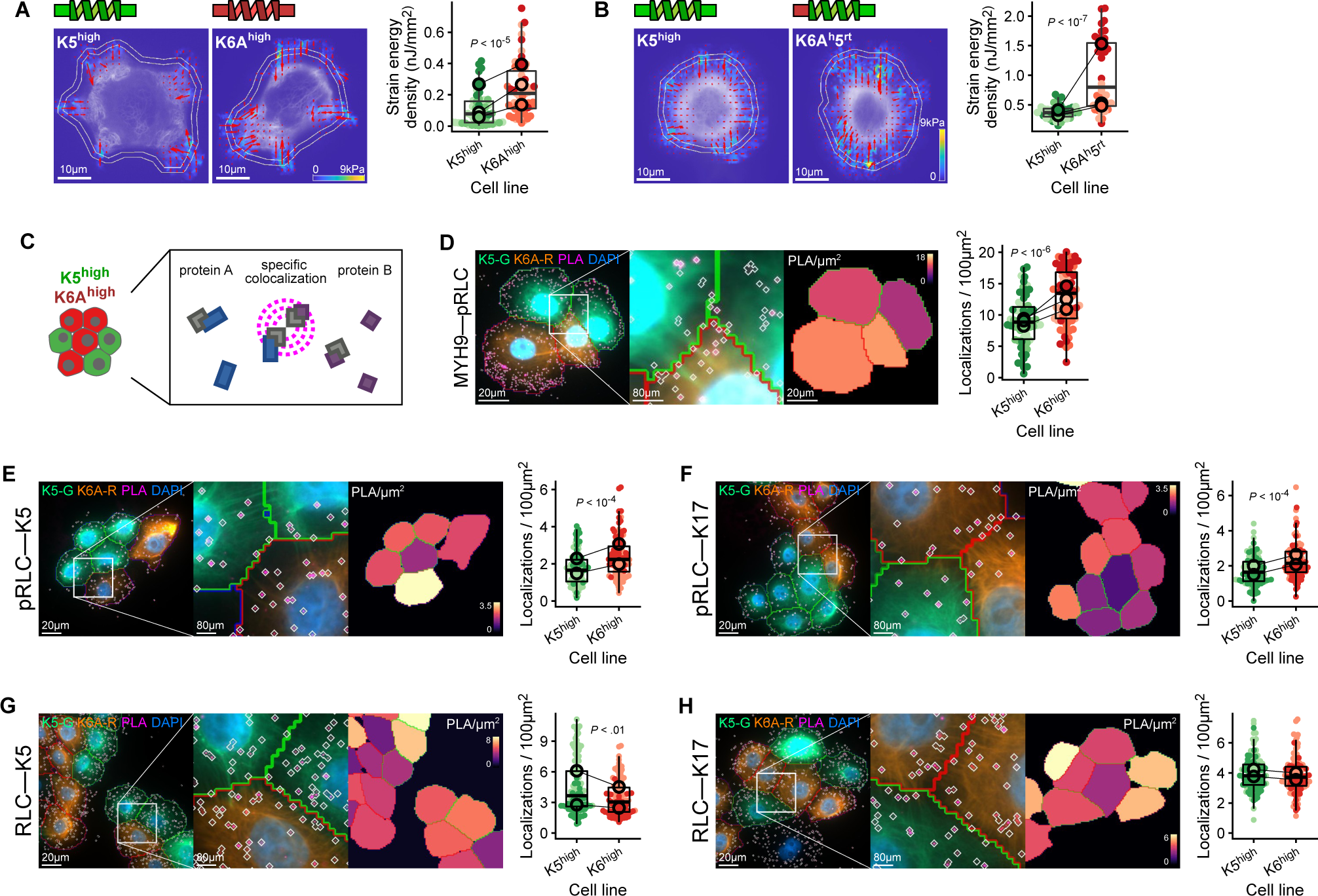
Wound-associated K6A increases cellular force generation by activating myosin. A. Traction force microscopy (TFM) of K5^high^ and K6A^high^ cells (see Methods). Left, reconstructed traction vectors (arrows) on a heatmap of force magnitude and a watermark of keratin filament images (partially captured by the TIRF field). Inner ring, cell border based on keratin images; Outer ring, expanded boundary for strain energy calculation (see also Figure S4C). Right, strain energy density in K5^high^ and K6A^high^ cells. Outlined dots, connected between groups, mean values of experimental replicates in separate batches of TFM substrates. *n* = 51-58 cells per group. B. TFM of cells expressing a chimeric keratin containing the head domain of K6A joined to the rod and tail of K5 (K6A^h^5^rt^) or full-length K5. See also Figure S4E. *n* = 40 cells per group. C. Schematic diagram of the proximity ligation assay (PLA). D. PLA detecting activated myosin by colocalization of phosphorylated myosin regulatory light chain (pRLC) and non-muscle myosin heavy chain (MYH9). Left, annotated images of K5^high^ (green outline) and K6A^high^ (red outline) cells. White diamonds, PLA detections (see Methods). Heatmap, PLA density (detections per 100 μm^2^). K5-G, K5-mNeonGreen; K6A-R, K6A-mRuby2. Right, PLA detection density in K5^high^ and K6A^high^ cells. Outlined dots, connected between groups, mean values of experimental replicates. *n* = 66-80 cells per group in 37 total images. E-F. PLA detecting activated myosin near keratin filaments by colocalization of pRLC and K5 (E) or K17 (F). E, *n* = 94-115 cells per group in 20 images. F, 99-108 cells per group in 23 images. G-H. PLA detecting total myosin near keratin filaments by colocalization of total myosin regulatory light chain (RLC) and K5 (G) or K17 (H). G, *n* = 117-119 cells per group in 19 images. H, 121-129 cells per group in 21 images.

Keratin filaments are not known to directly generate force. Rather, contractile forces are generated by myosin motor proteins linked to the actin cytoskeleton^46^. Non-muscle myosin activation occurs through phosphorylation of myosin regulatory light chain (pRLC) ^47,48^. To directly compare pRLC between K5^high^ and K6A^high^ cells, we employed a proximity ligation assay, which uses two primary antibodies raised in different species, oligonucleotide-conjugated secondary antibodies, and rolling-circle amplification to detect colocalization between two antigens (Figure 5C) ^49^. We probed for colocalization of myosin heavy chain (MYH9) and pRLC in a mosaic culture of K5^high^ and K6A^high^ keratinocytes. Consistent with the finding of increased contractility of K6A^high^ cells, we measured increased colocalization of pRLC with MYH9, indicating increased myosin activation, in K6A^high^ cells compared to K5^high^ cells (Figure 5D).

We further probed for colocalization of pRLC with keratin filaments using either antibodies against K5 or K17. In both cases, we detected increased colocalizations in K6A^high^ cells compared to K5^high^ cells (Figure 5E-F), suggesting that myosin motors in the neighborhood of keratin filaments are more frequently activated when the relative amount of K6A is increased. Importantly, the colocalization between total regulatory light chain and K5 or K17 was similar or decreased in K6A^high^ cells compared to K5^high^ cells (Figure 5G-H), underscoring that increased pRLC near keratin filaments must reflect a signaling phenomenon leading to myosin activation, not merely a differential physical coupling between keratins and the actomyosin machinery.

### Wound-associated K6A recruits non-filamentous vimentin to keratin filaments

We next asked how changing keratin filament composition could alter myosin activation. We hypothesized that keratin filaments might serve as a platform to spatially organize components of a signaling pathway leading to RLC phosphorylation. If this was the case, changing keratin filament composition might alter the affinity of keratin filaments for components of such a signaling pathway, increasing or decreasing signal transduction. Reliable identification of keratin filament binding partners has been difficult for two reasons. First, keratins are expressed in extraordinary abundance, representing about 30% of total protein in basal keratinocytes^50^, which increases the risk of identifying non-specific associations. Second, epidermal keratin filaments are highly insoluble; the harsh, high-salt buffers capable of completely solubilizing keratin filaments likely disrupt interactions between keratins and other proteins^51^ and denature antibodies used for immunoprecipitation^52^. While methods have been proposed to immunoprecipitate epidermal keratins using milder detergents^51,53^, it remains unclear if they reliably capture filamentous keratin versus soluble keratin monomers or smaller aggregates.

To overcome this challenge, we created keratinocyte cell lines expressing K5 or K6A fused to a promiscuous biotin ligase^54^. This allowed us to perform a proteomics screen comparing biotinylated proteins from each cell line^55^, identifying in a differential manner proteins preferentially associated with keratin filaments in K5^high^ cells versus K6A^high^ cells (Figure 6A-B). These hits were implicated in a variety of functional processes, including cytoskeletal organization (Figures 6C and S5A).

**Figure 6.**
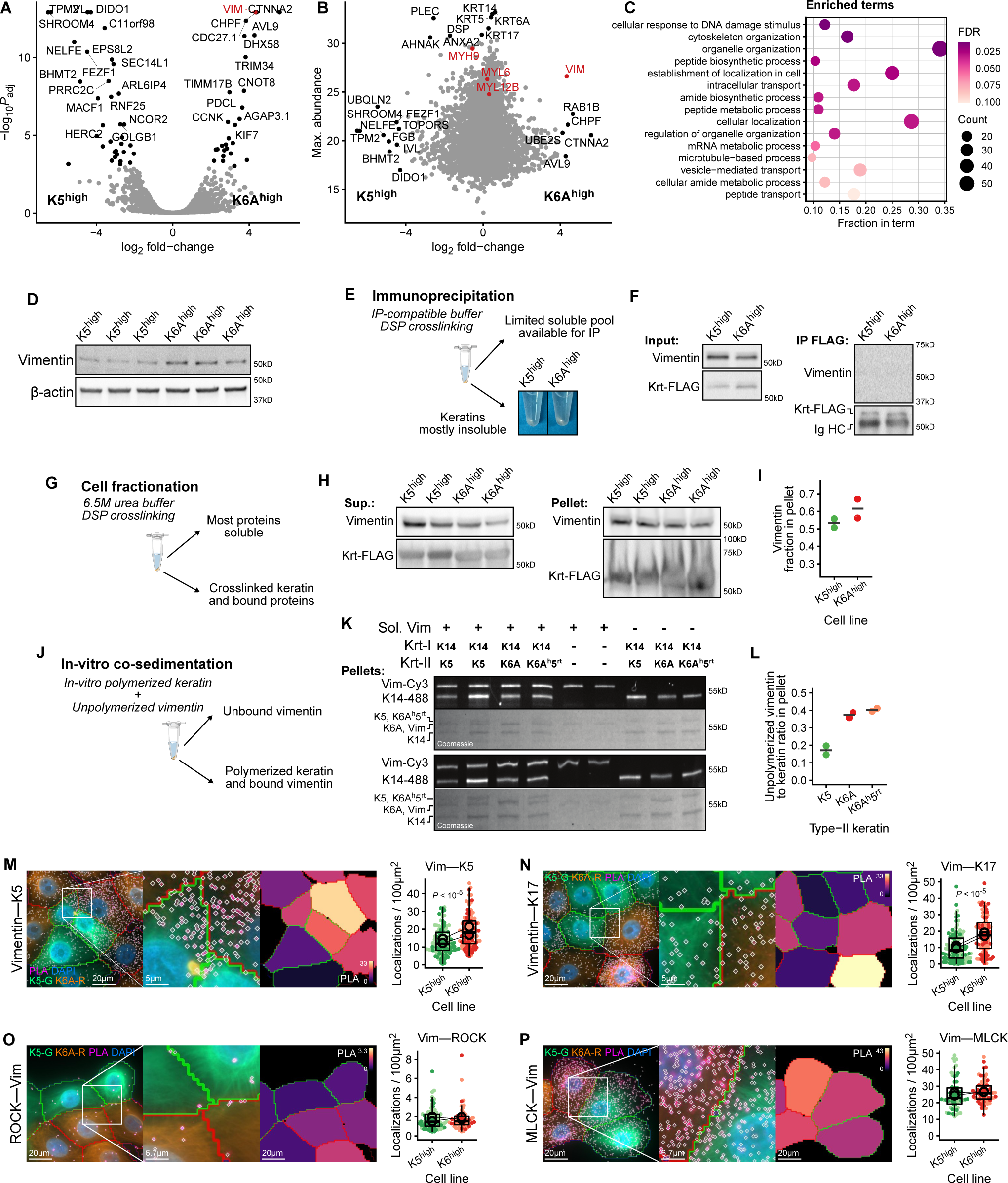
Wound-associated K6A recruits non-filamentous vimentin to keratin filaments. A-B. Proteins differentially associated with keratin filaments in K5^high^ cells versus K6A^high^ cells were identified by proximity biotinylation, streptavidin pulldown, and mass spectrometry (see Methods). A, Volcano plot highlighting differentially associated proteins. B, Differential association compared to the maximum protein abundance across all samples. *n* = 2 independent replicates per group. A. C. Most enriched Gene Ontology Biologic Process labels among proteins differentially associated with keratin filaments between K5^high^ and K6A^high^ cells, defined as greater than 3-fold differential association and *P*_adj_ < .05. See also Figure S5A. B. D. Western blot analysis of vimentin expression in K5^high^ and K6A^high^ cells. See also Figure S1E for additional blots with these samples; β-actin control repeated for reference. E-F. Keratin immunoprecipitation (IP) from K5^high^ and K6A^high^ cells expressing FLAG-tagged keratins. E, Schematic diagram of experiment with images of the large insoluble fraction remaining following lysis with an IP-compatible buffer (see Methods). F, Western blot analysis of the soluble fraction (Input) and IP with anti-FLAG beads. Krt-FLAG, tagged keratin; Ig HC, immunoglobulin heavy chain. G-I. Cell fractionation from K5^high^ and K6A^high^ cells expressing FLAG-tagged keratins. G, Schematic diagram of experiment. Following covalent crosslinking and lysis with a 6.5M urea buffer, lysates were separated into soluble and insoluble fractions (see Methods). H, Western blot analysis of the soluble supernatants (Sup.) and insoluble pellets. I, Quantification of the fraction of vimentin found in the pellet for each sample. J-L. Co-sedimentation analysis of in-vitro polymerized keratin filaments and unpolymerized vimentin. J, Schematic diagram of experiment. Fluorescently labelled K14 (K14-488) and either K5, K6A, or K6A^h^5^rt^ chimera were polymerized in vitro, mixed with unpolymerized fluorescently labeled vimentin (Vim-Cy3), and centrifuged at low speed to pellet intermediate filament polymers (see Methods). K, Fluorescence and Coomassie-stain images of gel electrophoresis of the pellets. Upper and lower gels represent independent experimental replicates. Note that each experimental replicate contains two each K5/K14 and vimentin control samples, which were averaged for quantification. L, Vimentin to keratin ratio in pellets, after subtracting the average amount of pelleted vimentin in the absence of keratin. M-N. Proximity ligation assay (PLA) detecting colocalization between vimentin and K5 (M) or K17 (N). Left, annotated images as in Figure 5D. Right, PLA detection density. Outlined dots, connected between groups, mean values of experimental replicates. M, *n* = 103-113 cells per group in 17 images. N, *n* = 84-105 cells per group in 18 images. O-P. PLA detecting colocalization between vimentin and ROCK (O) or MLCK (P). Note the different detection density ranges in (O) and (P). O, *n* = 87-121 cells per group in 20 images. P, *n* = 80-105 cells per group in 19 images.

To our great surprise, among the proteins with overall high keratin filament association, vimentin showed the highest differential association between K6A^high^ and K5^high^ cells (Figure 6A-B). Vimentin, a type-III intermediate filament protein, is the principal intermediate filament component in mesenchymal cells, and its expression at high levels in epithelial cells is associated with an epithelial to mesenchymal transition^56^. Based on classic two-dimensional gel electrophoresis^57^ and immunohistochemistry studies^35^ which failed to detect vimentin in the epidermis, it has typically been regarded as unimportant in that tissue. Yet these methods are relatively insensitive; more recently published proteomics datasets do detect vimentin in epidermis^58–60^, isolated primary keratinocytes^59^, and another keratinocyte cell line^61^, albeit at lower levels than keratin isoforms (Figure S5B). The role of epidermal vimentin in wound healing has also been unclear. While vimentin expression was increased at migration edges in some non-keratinocyte^62,63^ and cultured keratinocyte models^64,65^, vimentin expression was not reliably induced in skin wounds (Figures 1C and S5C), nor was vimentin clearly detectable by immunofluorescence at the migration edge in our epidermal culture wound model (Figure S5D). Furthermore, isoform-specific interactions between keratin filaments and vimentin have not previously been reported.

We first confirmed that our keratinocyte cell lines did express low levels of vimentin, with relatively increased vimentin in K6A^high^ cells compared to K5^high^ cells (Figure 6D). However, by immunofluorescence, vimentin filaments could only be detected in a very small fraction of keratinocytes (Figure S6). Even in this small minority of cells, vimentin filaments did not assemble into well-formed networks, but rather assembled into smaller aggregates or squiggles that did not align with keratin filaments (Figure S6) ^66^. The large majority of keratinocytes displayed no distinct vimentin structures at all, only a diffuse or speckled pattern, with no obvious difference between K5^high^ and K6A^high^ cells (Figure S6). Thus, we concluded that vimentin was present predominantly in non-filamentous form.

Unsurprisingly given the highly insoluble nature of keratin filaments, our attempts to capture the presumably transient keratin–vimentin interaction by co-immunoprecipitation from cell lysates were inconclusive, with very little keratin solubilized in immunoprecipitation-compatible lysis buffer (Figure 6E-F). Covalent crosslinking simply decreased keratin solubility even further, preventing full keratin solubilization even in 6.5M urea (Figure 6G-H). We took advantage of this seemingly inconvenient biophysical property to perform a cell fractionation experiment, reasoning that otherwise soluble keratin-interacting proteins would be pulled into the insoluble fraction (Figure 6G). We found that vimentin was indeed present in the keratin filament pellet, and although this fractionation may be only partially specific, there was a trend toward an increased proportion of vimentin in the insoluble fraction in K6A^high^ cells compared to K5^high^ cells (Figure 6H-I). Similarly, we were able to detect an interaction between in-vitro polymerized keratin filaments and non-polymerized vimentin using a co-sedimentation assay, and the amount of vimentin co-sedimentation was clearly increased with keratin filaments containing K6A or K6A^h^5^rt^ chimeras compared to K5 (Figures 6J-L and S7A).

Consistent with the proteomic screen and in vitro assay, we identified increased association between vimentin and keratins in K6A^high^ cells compared to K5^high^ cells by a proximity ligation assay, regardless of whether K5 or K17 was used as the bait (Figure 6M-N). Furthermore, the increased association between vimentin and keratin filaments in K6A^high^ cells was a population effect, not driven by a small number of outlier cells which might represent those with vimentin filaments (Figures 6M-N and S6). Together, these data led us to the conclusion that a non-filamentous pool of vimentin interacts with keratin filaments, preferentially with the wound-associated keratin isoform K6A.

### Non-filamentous vimentin shuttles myosin-activating kinases to K6A-contianing keratin filaments

How could association between non-filamentous vimentin and keratin promote actomyosin contractility through phosphorylation of myosin regulatory light chain? We considered the possibility that non-filamentous vimentin operates as a shuttle for kinases, bringing them into the proximity of keratin filaments, with a bias toward filaments with high K6A content. There is precedence for such a role, as non-filamentous vimentin has been implicated as a transport adaptor for MAP kinase in neurons^67^.

Furthermore, vimentin was previously identified as a phosphorylation target for ROCK, triggering filament disassembly^68^, and implying at least transient ROCK–vimentin binding. ROCK also targets RLC^69^. While we detected only infrequent colocalization between vimentin and ROCK by proximity ligation (Figure 6O), we did detect frequent colocalization between vimentin and MLCK (Figure 6P), another RLC activator^47^. Neither ROCK nor MLCK were differentially associated with vimentin in K5^high^ compared to K6A^high^ cells (Figure 6O-P). However, once differentially shuttled to keratin filaments by vimentin, these kinases would be positioned near other keratin-associated proteins. Myosin subunits were previously shown to bind keratins^33^, a finding recapitulated in our proteomics screen regardless of keratin isoform composition (Figure 6B). Thus, increasing the amount of K6A in keratin filaments would increase recruitment of regulatory kinases to activate myosin (Figure 7A).

**Figure 7.**
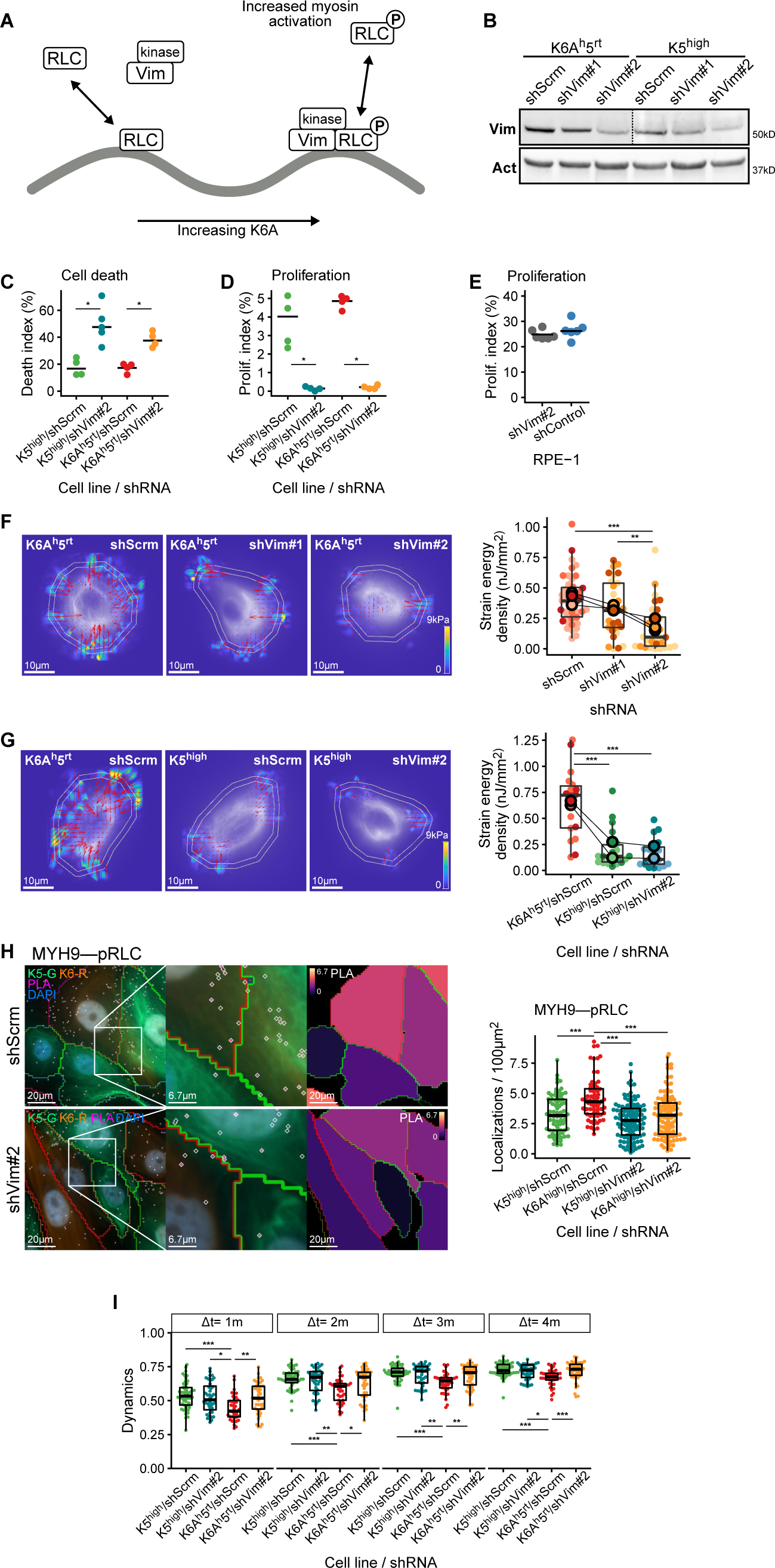
Vimentin shuttles regulatory kinases to activate myosin near K6A-containing filaments. A. Schematic diagram of the proposed model. Non-filamentous vimentin (Vim) shuttles regulatory kinases preferentially to K6A-containing filaments, where they phosphorylate myosin regulatory light chain. B. Western blot analysis of vimentin knockdown by shRNA (shVim#1; shVim#2; scrambled control, shScrm) in keratinocytes expressing a chimeric keratin containing the head domain of K6A joined to the rod and tail of K5 (K6A^h^5^rt^) or full-length K5 (K5^high^). Dashed line separates different brightness adjustments for different baseline vimentin levels. See also Figure S7B for an alternate display. C. Death of K5^high^ and K6A^h^5^rt^ cells upon expression of shVim#2 versus shScrm. Cell death index, percentage of nuclei with membrane-impermeable DNA dye uptake (see Methods). *, *P* < .05. *n* = 4-5 samples with 112-162 total cells per group. D. Proliferation of K5^high^ and K6A^h^5^rt^ cells upon expression of shVim#2 versus shScrm. Proliferation index, percentage of cells with EdU uptake (see Methods). *, *P* < .05. *n* = 4 samples with 2,897-6,013 total cells per group. E. Proliferation of RPE-1 retinal epithelial cells upon expression of shVim#2 versus shScrm. *n* = 6 samples with 1,853-1,930 total cells per group. F. Traction force microscopy (TFM) of K6A^h^5^rt^ cells upon expression of shVim#1, shVim#2, or shScrm. Left, reconstructed traction force vectors (arrows) on a heatmap of force magnitude and a watermark of keratin filament images. Inner ring, cell border based on keratin images. Outer ring, expanded boundary used for strain energy calculation. Right, comparison of strain energy densities between treatment groups. Outlined dots, connected between groups, mean values of experimental replicates in separate batches of TFM substrates. ***, *P* < 10^-3^; **, *P* < .01. *n* = 29-43 cells per group. G. TFM of K6A^h^5^rt^ or K5^high^ cells upon expression of shVim#2, or shScrm. Annotations as in (F). *n* = 18-21 cells per group. H. PLA detecting myosin activation by colocalization of phosphorylated myosin regulatory light chain (pRLC) and non-muscle myosin heavy chain (MYH9) in K6A^high^ and K5^high^ cells upon expression of shVim#2 or shScrm. ***, *P* < 10^-3^. *n* = 67-113 cells per group in 16-24 images per shRNA condition. I. Keratin filament network dynamics in K6A^h^5^rt^ cells and K5^high^ cells upon expression of shVim#2 or shScrm. See also Figure S7C for filament images, segmentations, and dynamics score maps. ***, *P* < 10^-3^; **, *P* < .01; *, *P* < .05. *n* = 39-49 cells per group.

This model implies a central role for vimentin mediating the activation of myosin and increased force generation with increased expression of wound-associated K6A. To test this, we knocked down vimentin in K5^high^ and K6A^h^5^rt^ keratinocyte cell lines using lentiviral shRNA vectors (Figures 7B and S7B). Vimentin knockdown was poorly tolerated by these keratinocytes, with decreased proliferation and increased cell death within one week of shRNA expression (Figure 7C-D), consistent with previously reported findings in primary human keratinocytes^70,71^. Absence of proliferation defects in a non-keratinocyte cell line decreased our concern for off-target effects (Figure 7E). Thus, we were unable to establish stable vimentin-depleted cell lines. However, by transient shRNA expression, we could perform traction force microscopy on vimentin knockdown keratinocytes and confirmed that vimentin depletion decreased traction force generation in K6A^h^5^rt^ cells (Figure 7F). Vimentin depletion only slightly decreased traction force generation in K5^high^ cells, likely reflecting the lower baseline contractility of K5^high^ cells compared to K6A^h^5^rt^ cells (Figure 7G). Consistent with this, vimentin knockdown decreased myosin activation in K6A^high^ cells to a level similar to that of K5^high^ cells (Figure 7H). Furthermore, vimentin knockdown reversed the effect of K6A on keratin filament dynamics. Compared to a scrambled shRNA control, vimentin shRNA increased filament dynamics in K6A^h^5^rt^ cells nearly to the level seen in K5^high^ cells (Figures 7I and S7C). Filament dynamics were little changed by vimentin knockdown in K5^high^ cells (Figures 7I and S7C). Altogether, these results show that increased traction force generation and decreased filament dynamics caused by wound-associated K6A both depend on non-filamentous vimentin acting as a shuttle for kinases.

## Discussion

Our data reveal an unexpected role for keratin filaments as isoform-tuned signaling scaffolds in addition to their canonical role as mechanical scaffolds. Increasing expression of wound-associated K6A supports epidermal remodeling not by altering the cell’s mechanical capacity for force resistance, but by organizing molecular signals that activate force generation to drive migration. This signal transduction cascade relies on an isoform-preferential interaction between the intrinsically disordered K6A head domain and non-filamentous vimentin shuttling myosin activating kinases. As the amount of K6A incorporated into keratin filaments increases, more kinase is shuttled to keratin filaments where it phosphorylates nearby RLC, leading to myosin activation.

Signaling functions of keratin filaments have long been postulated, ^72–74^ but how such signaling might operate in the context of different keratin isoform mixtures has not been clear. Most keratin signaling proposals rely on knock-out models or filament-disrupting mutants which are confounded by mechanical defects^18,75–83^. In one notable example, and in seeming disagreement with the data presented here, keratinocytes from a Krt6a/b-null mouse migrated faster than wild-type controls, purportedly through a Src- and myosin-mediated mechanism^18,31–33^. Yet these studies also noted broad disruption of cell-cell adhesion, ^18,31–33^ and two other mouse models with varying degrees of K6 depletion but apparently preserved mechanical integrity demonstrated unchanged or decreased keratinocyte migratory ability^17,21^. We suggest as a plausible unifying interpretation that increased keratinocyte migration in the Krt6a/b-null mouse results from general loss of keratin mechanical scaffold function and cell-cell adhesion, rather than isoform-specific effects of K6 per se.

A key finding of our study, uncovered only by preserving the keratin filament network and mechanical scaffold, is that keratin isoform-specific interactions allow shifts in the isoform mixture to tune molecular signals. In contrast, in the case of the Krt6a/b-null mouse, in vitro experiments suggested that keratin interactions with Src and myosin were not necessarily isoform-specific. ^32,33^ In the case of myosin, these finding are entirely consistent with our proximity proteomics data, although we did not detect Src in our dataset. More generally, we argue that filament-network-disrupting models cannot establish the isoform-specificity of keratin interactions, and without isoform-specific interactions, it is difficult to explain why the keratin isoform mixture shifts in contexts such as wound healing.

The pathway proposed here establishes a functional coupling between the cellular machinery of force resistance and force generation precisely during the context where the stability–plasticity balance must be finely tuned. During wound healing, the epidermis remains subject to mechanical stress, both from the usual external insults and directly resulting from tissue remodeling, so disassembling the keratin mechanical scaffold to increase tissue plasticity is an untenable strategy. Shifting the keratin isoform mixture therefore represents an elegant mechanism to maintain the mechanical capacity for force resistance while modulating non-mechanical functions through cellular signals.

In a seeming paradox, we observed that elevated K6A expression increases force generation yet decreases intracellular movement of keratin filaments. This may simply reflect impairment of keratin filament movement by heightened cellular rigidity or subtly increased formation of actomyosin stress fibers. However, an intriguing alternative hypothesis is that non-filamentous vimentin acts not only as a kinase shuttle, but also as a cytoplasmic glue, sticking keratin filaments to other cellular components such as stress fibers^84,85^, thus decreasing K6A-enriched keratin filament dynamics under a given force. Wound-associated keratins might then function both to activate actomyosin contractility and to stabilize otherwise brittle actin structures^86^. Expansive additional investigation will be needed to evaluate these possibilities.

Our results also highlight an underappreciated role of vimentin in epithelial tissues. In contexts such as development and cancer metastasis, vimentin expression is a marker of epithelial to mesenchymal transition (EMT) ^87^, but in stable epithelia vimentin has been thought to be irrelevant. Wound healing can be considered a reversible forme fruste of EMT^87^, and defective wound healing is among the most prominent phenotypes of a vimentin knock-out mouse^88^. Although this has been attributed to defects in dermal fibroblasts^89^, our results suggest that the keratinocytes themselves may significantly contribute to the phenotype because of their dependence on vimentin as a shuttle for molecular signals. Furthermore, our finding that vimentin knockdown compromises keratinocyte growth in culture despite the absence of vimentin filaments, consistent with previously reported findings^70,71^, underscores the importance of non-filamentous vimentin^90^ for cellular homeostasis, even in epithelial cells.

Despite clear in vitro biochemical and in situ cell biological evidence of a preferential interaction between vimentin and keratin filaments enriched for the K6A head domain, the precise structural nature of the interaction remains to be determined. Recent reports that intrinsically disordered intermediate filament head and tail domains interact through labile cross-β-strand polymerization suggest one intriguing possibility. ^91,92^ Post-translational modifications could also be involved. ^93^ The high degree of head and tail sequence variability between keratin isoforms establishes a thus far undiscussed potential for cell regulation through complex interaction networks between the intermediate filament cytoskeleton and other cytoplasmic proteins and subcellular structures. Such interactions could allow intermediate filaments to spatiotemporally organize a variety of biochemical processes, akin to the signaling pathway leading to actomyosin activation by wound-associated K6A identified here.

Different keratin isoform mixtures serve as reliable markers of fundamental epithelial states, distinguishing simple versus stratified epithelia, stem cell versus differentiated cell compartments, and, as we have focused our attention in this study, stable versus actively remodeling tissues. Our data suggest these isoforms serve not only to mark different epithelial states, but in fact establish those states through organization of signaling pathways or transcriptional networks. Thus, this study sets the stage for investigations of the diversity of intermediate filament interactions as a new regulatory motif in the adaptive control of cell function.

### Limitations of the study

Although we detected no difference in mechanical integrity between K5^high^ and K6A^high^ cells, it remains possible that different keratin isoforms may influence the material properties of filaments, the kinetics of subunit exchange, or the formation of desmosomes or hemidesmosomes, including in potentially subtle ways via feedback loops related to actomyosin contractility. Dedicated studies will be needed to address these questions. Additionally, the full mechanistic link between actomyosin activation and increased wound closure is likely complex and is not comprehensively elucidated here. Given that K6A^high^ cells possessed a transient migration advantage in two-dimensional culture but a persistent advantage in three-dimensional epidermal culture, we speculate that environmental cues significantly modulate this behavior. Future studies will explore this interaction.

## Acknowledgements

We thank Kim Reed and other members of the Danuser laboratory for technical support and helpful discussion; Jerry Shay, Kimberly Batten, Richard Wang and Travis Vandergriff for reagents and advice; John Shelton, Diana Wigginton, and Cameron Perry from the UT Southwestern Histo Pathology Core for histology services and technical support; the Moody Foundation Flow Cytometry Facility of the Children’s Medical Center Research Institute at UT Southwestern; the UT Southwestern Flow and Mass Cytometry Facility; the UT Southwestern Electron Microscopy Core Facility, supported in part by the National Institutes of Health (S10OD021685); the UT Southwestern Proteomics Core Facility; and the BioHPC computing facility at UT Southwestern. B.A.N. is supported by a Career Development Award from the Dermatology Foundation, the National Institutes of Health (T32AR065969), the UT Southwestern Physician Scientist Training Program, and the Foundation for Ichthyosis and Related Skin Types. Research in the Danuser lab is supported by grants from the National Institute of General Medical Sciences (R35GM136428 and RM1GM145399 for dissemination of tools). Research in the Medalia lab is supported by a grant from the Swiss National Science Foundation (SNSF 310030_207453).

## Methods

### Cell culture

Human keratinocyte Ker-CT cells (ATCC #CRL-4048) were a kind gift from Dr. Jerry Shay (UT Southwestern Medical Center). This cell line was developed from human foreskin keratinocytes through expression of HTERT and CDK4. ^28,29^ Unless otherwise specified, keratinocytes were cultured on tissue-culture treated plastic coated with bovine type I collagen (PureCol, Advanced Biomatrix #5005) by pre-incubating dishes in a 100 μg/mL collagen solution for 30 minutes. Cells were maintained in Keratinocyte serum-free medium (K-SFM; Gibco #17005042) with the included 5 ng/mL human recombinant epidermal growth factor and 50 mg/mL bovine pituitary extract supplements according to the manufacturer’s instructions in a humidified incubator at 37 °C and 5% CO_2_. K-SFM contains a low calcium concentration (about 0.12mM according to the manufacturer) which supports keratinocyte growth without inducing differentiation, but which also limits desmosome formation^94^. To induce desmosome formation and support differentiation for certain experiments, we developed a keratinocyte differentiation medium based on a simplification of the classic Rheinwald and Green formula^95,96^. This medium was composed of fully-supplemented K-SFM mixed in equal parts with DMEM/F12 (Gibco #11320033) and additionally supplemented with 2% fetal bovine serum (FBS; Sigma F0926). Based on an estimated calcium concentration of 3.75mM in FBS^97^, the calcium concentration of the differentiation medium was 0.65mM. As indicated below, 0.5mM additional calcium chloride, yielding a final concentration of 1.15mM, was added for some experiments. HEK293 cells (a gift from H. Choe) used for generation of lentiviral particles were maintained in DMEM (Gibco #11995-065) supplemented with 10% FBS and antibiotic–antimycotic reagent (Gibco #15240-062) in a humidified incubator at 37 °C and 5% CO_2_. hTERT RPE-1 cells were obtained from ATCC (CRL-4000) and maintained in equal parts DMEM/F12 supplemented with 10% FBS in a humidified incubator at 37 °C and 5% CO_2_. All cell lines were periodically tested for mycoplasma using a PCR-based Genlantis Mycoscope Detection Kit (MY01100).

### Plasmids

Primer and synthetic DNA sequences (Integrated DNA Technologies) used to assemble the following constructs are provided in Table S1.

Keratin sequences were cloned by polymerase chain reaction (PCR) from pBabe-RFP1-KRT5-hygro (Addgene #58493) and pDONR22-KRT6A (DNASU clone HsCD00039474) and inserted into pLVX-IRES-Puro or pLVX-IRES-Neo lentiviral expression vectors (Clontech) with C-terminal fluorescent tags using seamless cloning (HiFi DNA Assembly Master Mix, New England Biolabs). The pLVX-KRT5-mNG-IRES-Puro construct includes an mNeonGreen (Allele Biotechnology) ^98^ sequence in-frame with the KRT5 sequence separated by a short peptide linker (DPAFLY). Similarly, the pLVX-KRT6A-mRb-IRES-Neo construct includes an mRuby2 (Addgene #40260) ^99^ sequence in-frame with the KRT6A sequence separated by the same linker.

Keratin head/tail domain chimera constructs were assembled using seamless cloning into pLVX-IRES-Puro, with keratin domains cloned from the above pLVX-KRT5-mNG-IRES-Puro and pLVX-KRT6A-mRb-IRES-Neo constructs by PCR. Keratin domains were defined based on NCBI Reference Sequence annotations (NP_000415.2, NP_005545.1). Specifically, the K5 head and tail domains included amino acids 1-167 and 478-590 respectively; the K6A head and tail domains included amino acids 1-162 and 473-564 respectively.

For proximity biotin labeling experiments, the sequence of miniTurbo, a small promiscuous biotin ligase^54^, was synthesized with a C-terminal Flag tag and flanking XbaI and BamHI restriction sites. This fragment was then inserted into pLVX-IRES-Puro using the XbaI and BamHI sites. Finally, the KRT5 and KRT6A sequences were amplified by PCR from the above pLVX-KRT5-mNG-IRES-Puro and pLVX-KRT6A-mRb-IRES-Neo constructs and inserted in-frame and N-terminal to the biotin ligase with a short peptide linker (DPAFSR) using seamless cloning. To create a marker construct for these experiments, EGFP from pEGFP-C1 (Clontech) was inserted into pLVX-IRES-Puro using the SnaBI and BamHI restriction sites.

To create short-tagged keratin constructs without linked fluorescent proteins, the Q5 Site-Directed Mutagenesis Kit (NEB) was used to delete the miniTurbo sequence from the above pLVX-KRT5-miniTurbo-FLAG-IRES-Puro and pLVX-KRT6A-miniTurbo-FLAG-IRES-Puro constructs, leaving the FLAG joined in-frame to the C-terminal end of the keratin sequence. Seamless cloning was then used to replace the puromycin resistance cassette after the internal ribosome entry sequence with an mNeonGreen sequence, allowing for expression of a non-linked fluorescent marker from the same plasmid.

For shRNA experiments, pLKO.1-Scrambled (Addgene #136035) ^100^ was modified to express H2B-mRuby3 for visual identification of transduced cells based on a nuclear fluorescence signal^101^. The H2B-mRuby3 sequence was synthesized and inserted into pLKO.1 in place of the existing PuroR sequence using BamHI and KpnI restriction sites. Vimentin targeting sequences for shRNA constructs were selected from the Genetic Perturbation Platform Web Portal (Broad Institute; https://portals.broadinstitute.org/gpp/). Complementary primers were synthesized and annealed, then cloned into pLKO1-H2B-mRuby3 using AgeI and EcoRI restriction sites. The clones selected for shVim#1 (TRCN0000029120) and shVim#2 (TRCN0000029119) have, respectively, moderate and high predicted knockdown performance.

For in-vitro expression and fluorescent labeling of vimentin, the sequence encoding human full-length vimentin (amino acids 1-466) was modified to remove its single internal cysteine residue (C328A) and add a GGC tripeptide to the C-terminal end. Then the sequences encoding vimentin^C328A^-GGC, wild-type vimentin, KRT5, KRT6A, and K6A^h^5^rt^ were cloned into pEt24d(+) (EMD Biosciences) by PCR.

### Lentivirus transduction and cell line selection

Lentiviral particles were generated using the pLVX system (Clontech) with packaging vectors psPAX2 and pMD2.G (Addgene plasmids #12260 and #12259). HEK293 cells were transfected with expression and packing plasmids following standard calcium phosphate or polyethylenimine (Polysciences #23966) protocols. Supernatant was collected two days after transfection, filtered through 0.45-μm mixed cellulose esters membrane syringe filters (Fisher Scientific #09-720-005) and incubated on target cells overnight.

After multiple days of culture and cell expansion, cells expressing the relevant fluorescent marker were collected on a FACS Aria II SORP flow cytometer equipped with 405 nm, 488 nm, 561 nm, and 633 nm lasers (Beckton Dickinson) and returned to culture.

### Western blotting

Cells were grown to confluence and switched to differentiation medium 24 hours prior to sample collection. Because keratin filaments are highly insoluble, a high-ionic-strength urea buffer was used in place of a standard lysis buffer^51^. The 6.5 M urea lysis buffer also contained 50 mM tris, 1 mM EGTA, 2 mM DTT, 50 mM sodium fluoride, 1 mM sodium vanadate, and Halt Protease Inhibitor Cocktail (ThermoFisher #78430), and was adjusted to pH 7.5. After washing twice with PBS, cells were collected into urea lysis buffer using a cell scraper, then incubated at 4 °C for 20 minutes on a shaker. The lysates were then sonicated using a Fisherbrand Model 505 Sonic Dismembrator (ThermoFisher) twice for 10 seconds at 20% power with a 10 second intervening rest. Any remaining insoluble material, which was typically negligible, was pelleted by centrifugation at 16.1×10^3^ *g* for 20 minutes at 4 °C, and the supernatant was transferred to a clean tube. Protein concentrations were measured using a BCA assay (ThermoFisher #23228). Laemmli sample buffer (BioRad #161-0747) with 2-mercaptoethanol (ThermoFisher #BP176) was added to the samples, which were then heated to 95 °C for 5 minutes.

Samples were run on 4-20% Mini-PROTEAN pre-cast gels (Biorad) and transferred to nitrocellulose membranes (ThermoFisher #88018). Once membranes were blocked with 5% bovine serum albumin (BSA; Equitech-Bio BAH65-0500) in Tris buffered saline (TBS) with 0.1% Tween 20 (TBST; ThermoFisher BP337) for 1 hour at room temperature, they were incubated in primary antibodies overnight at 4 °C (Table S2). Membranes were washed with TBST and probed with secondary antibodies conjugated with horseradish peroxidase (Table S2). Once they were washed again, bands were detected through enhanced chemiluminescence using freshly prepared substrate solution based on 100 mM Tris (ThermoFisher BP152) adjusted to pH 8.5 with hydrochloric acid (ThermoFisher A144S-212), 1.2 mM luminol (Sigma A8511), 2.5 mM p-coumaric acid (Sigma C9008) and 0.04% hydrogen peroxide (w/v, Alpha Aesar L14000) on a Syngene G:BOX imager equipped with a Synoptics 4.2-megapixel camera and controlled with Syngene Genesys software.

### Quantitative reverse transcription PCR

Keratinocytes were grown to confluence in 6-well tissue culture plates. For comparison between cell lines (Figure S1G), medium was switched from K-SFM to differentiation medium 24 hours before sample collection. For scratch wound experiments (Figure 1F), culture medium was switched from K-SFM to differentiation medium, then treatment samples were scratched with a 1000 mL pipette tip in a cross-hatch pattern. The scratch procedure was repeated every other day for 10 days. Samples were collected one day following the last scratch procedure. RNA was isolated from the samples using RNeasy spin column kits (Qiagen #74004) and reverse transcription was performed using SuperScript IV First-Strand Synthesis System with oligo(dT)_20_ primers (Invitrogen #18091050). Real-time PCR using a C1000 thermal cycler with a CFX96 optical reaction module (Bio-Rad), iTaq Universal SYBER Green Supermix (Bio-Rad), and transcript-specific primers (Table S1) was performed to quantify transcript levels. The 2^−ΔΔCT^ method^102^ was used to compare each individual keratin to the mean level of all keratins measured.

### Keratinocyte monolayer fragmentation assay

We adopted the commonly used dispase-based fragmentation assay to verify the ability of our keratinocyte cell lines to form stable monolayer barriers^103^. Keratinocytes were seeded into collagen-coated 12-well tissue culture plates and grown to confluence. Upon reaching confluence, culture medium was changed from K-SFM to differentiation medium with additional 0.5 mM calcium chloride. After an additional 48-hour incubation, the monolayers were washed twice with PBS then incubated in dispase solution (5 units/mL in HBSS, Stemcell Technologies #07913) for 80 minutes at 37 °C to detach the monolayers from the plate. The detached monolayers were then exposed to mechanical shear stress by slow passage through a 1000μL pipette tip 10 times. Each plate was then imaged twice with an Apple iPhone 13 mini with gentle rocking between the image pairs to displace the monolayer fragments.

Monolayer fragments were manually annotated and counted. If a different number of fragments was counted in each of the image pairs for a given well, the higher value was used.

### Type-II wound-associated keratin cluster heterozygous deletion

The human genome contains three type-II wound-associated keratin genes, KRT6A, KRT6B, and KRT6C, located in series with no other intervening genes in a 48 kilobase region on chromosome 12. We adopted a two-stage strategy of loxP cassette insertion followed by Cre recombination to attempt deletion of the entire genomic cluster. First, we used CRISPR-Cas9 with homology directed repair to insert selection cassettes at sites flanking the keratin gene cluster. Each cassette contained a fluorescent protein expression sequence (either mNeonGreen or mCherry) and a loxP site, designed such that if the cassettes were correctly positioned and oriented, Cre-mediated recombination would remove the keratin gene cluster along with the fluorescent markers. Each cassette was synthesized and inserted into a tia1l self-targeting carrier plasmid^104^ by seamless cloning. Guides were synthesized and inserted into the pX333 system (Addgene #64073), which expresses tandem sgRNAs as well as Cas9^105^. The three plasmids were transfected together into keratinocytes using Lipofectamine LTX with PLUS reagent (Invitrogen #A12621) according to the manufacturer’s protocol. After 7 days in culture to allow cassette insertion and cell expansion, cells expressing both fluorescent markers (about 1% of the total population) were collected by FACS and returned to culture. After an additional 20 days in culture to ensure stable integration of the cassettes, cells were again sorted for expression of both fluorescence markers. To finally excise the keratin cluster, we transduced the cells with Ad5-CMV-Cre adenoviral vector (purchased from Baylor College of Medicine Gene Vector Core). After 7 days of culture, cells lacking both fluorescent markers, indicating successful deletion of the keratin cluster, were collected by FACS, and clonal lines were created by limiting dilution. We were unable to isolate clones completely lacking K6. However, we did isolate a clone with K6 protein decreased by approximately 50% (Figure S1D). We isolated genomic DNA from this clone using a Qiagen DNEasy kit and used PCR to amplify a region from the type-II wound-associated keratin cluster as well as a region formed by the predicted deletion.

Both bands were present, indicating a heterozygous clone (Figure S1D). This cell line was used as a positive control for the keratinocyte monolayer fragmentation assay (Figure S1C).

### Keratin immunoprecipitation

K5^high^ and K6A^high^ cell lines expressing FLAG-tagged keratin constructs were grown to confluence and switched to differentiation medium 24 hours prior to sample collection. Cells were washed with PBS, then incubated in 1 mM dithiobis(succinimidyl propionate) (DSP, ThermoFisher #22585) on ice for 1-hour to covalently crosslink interacting proteins. The reaction was quenched by incubating in 25 mM Tris, pH 7.5 on ice for 15-minutes. Cells were then washed with PBS and lysed in an immunoprecipitation buffer reported to partially solubilize keratins (40 mM HEPES, 120 mM sodium chloride, 1 mM EDTA, 1% Triton X-100, 2% Empigen BB [Sigma-Aldrich #30326], and protease inhibitors [Halt Protease and Phosphatase Inhibitor, Thermo], pH 7.5). ^51,53^ After incubation on a shaker at 4 °C for 20-minutes, lysates were sonicated using a Fisherbrand Model 505 Sonic Dismembrator (ThermoFisher) twice for 10 seconds at 20% power with a 10 second intervening rest. The lysates were then centrifuged at 16.1×10^3^ *g* for 20 minutes at 4 °C, producing a substantial pellet (Figure 6E). The supernatant was transferred to a clean tube and protein concentrations were measured using a BCA assay (ThermoFisher #23228).

For each sample, 150 µL of anti-FLAG M2 magnetic beads slurry (Millipore Sigma M8823) was prepared in a microcentrifuge tube by washing three times with tris-buffered saline (TBS; 50 mM Tris, 150 mM sodium chloride, pH 7.4) and resuspending in 50 µL of TBS. Lysate with 500 µg of protein was added to the beads and incubated with rotation at 4 °C for 2-hours. Remaining lysate (input) was combined with Laemmli sample buffer (BioRad #161-0747) with 2-mercaptoethanol (ThermoFisher #BP176). Following incubation, the beads were washed three times with TBS, then bound protein was eluted directly in Laemmli sample buffer with 2-mercaptoethanol. All samples were denatured at 95 °C for 5 minutes and analyzed by Western blot.

### Cell fractionation for crosslinked keratin filaments

K5^high^ and K6A^high^ cell lines expressing FLAG-tagged keratin constructs were grown to confluence and switched to differentiation medium 24 hours prior to sample collection. Cells were washed with PBS, then incubated in 1 mM dithiobis(succinimidyl propionate) (DSP, ThermoFisher #22585) on ice for 1-hour to covalently crosslink interacting proteins. The reaction was quenched by incubating in 25 mM Tris, pH 7.5 on ice for 15-minutes. Cells were then washed with PBS and lysed in 6.5 M urea buffer (50 mM tris, 1 mM EGTA, 50 mM sodium fluoride, 1 mM sodium vanadate, and Halt Protease Inhibitor Cocktail [ThermoFisher #78430], pH 7.5; the reducing agent DTT, which would cleave the DSP crosslinker, was excluded from the buffer). ^51^ After incubation on a shaker at 4 °C for 20-minutes, lysates were sonicated using a Fisherbrand Model 505 Sonic Dismembrator (ThermoFisher) twice for 10 seconds at 20% power with a 10 second intervening rest. The lysates were then centrifuged at 16.1×10^3^ *g* for 20 minutes at 4 °C. Unlike keratinocyte lysates prepared without crosslinking in 6.5 M urea buffer as described above in “Western blotting,” the crosslinking procedure resulted in a noticeable pellet. The supernatants, representing the soluble fraction, were transferred into clean tubes and combined with Laemmli sample buffer (BioRad #161-0747) with 2-mercaptoethanol (ThermoFisher #BP176). The pellets, representing the insoluble fraction, were washed in lysis buffer then dissolved directly in Laemmli sample buffer with 2-mercaptoethanol. All samples were denatured at 95 °C for 5 minutes and analyzed by Western blot.

### In-vitro purification and reconstitution of intermediate filaments

Keratin or vimentin constructs in pEt24d(+) were expressed in Rosetta2 pLysS bacteria as inclusion bodies. The proteins were purified in multiple steps consisting of isolation of the inclusion bodies, solubilization in guanidinium chloride (GuHCl) for vimentin or in urea for keratins, and purification by size-exclusion chromatography. Briefly, the cell extract obtained after sonication of the frozen bacteria pellet in a buffer containing 50 mM Tris HCl pH 8, 200 mM NaCl, 25% Glycerol, 1 mM EDTA, 10 mg/ml lysozyme, 20 mM MgCl2, 8 ug/mL DNase 1, 40 ug/mL RNase A, 1% NP40, 1% Deoxycholic Acid sodium, and protease inhibitors (Roche cOmplete Protease Inhibitor Cocktail), was centrifuged for 30min at 12,000×*g*, at 4 °C. The inclusion bodies pellet was washed 3 times using cycles of resuspension/centrifugation (30 min at 12,000 g, at 4 °C) in a buffer containing 10 mM Tris HCl pH 8, 0.5% Triton X-100, 5 mM EDTA, 1.5 mM DTT, protease inhibitors (Roche cOmplete Protease Inhibitor Cocktail). As a last step the inclusion bodies pellet was washed in a buffer containing 10 mM Tris HCl pH 8, 1 mM EDTA, 1.5 mM DTT, and protease inhibitors (Roche cOmplete Protease Inhibitor Cocktail).^106^

The purified inclusion bodies pellets were further processed to produce renatured proteins and filaments. For vimentin, the inclusion bodies pellet was solubilized with 6M GuHCl in 10 mM TrisHCl pH 7.5, and clarified for 30 min at 10,000×*g* at 4 °C, and the vimentin-containing supernatant was collected. A size-exclusion chromatography using Superdex 200 Increase 10/300 GL (Cytiva) was performed and protein purity was checked using SDS-PAGE. The protein concentration was adjusted to 0.2-0.4 mg/ml, and vimentin was renatured by serial dialysis of 30 min at 22 °C using buffers of decreasing concentration of GuHCl, followed by an overnight dialysis step at 4 °C in the GuHCl free buffer. The dialysis buffers were 5 mM Tris HCl pH 7.5, with 1 mM EDTA, 0.1 mM EGTA, 1 mM DTT, containing 4, 2, 0 M GuHCl. For filament reconstitution, the protein solution was dialyzed against a high-salt buffer containing 10 mM Tris HCl pH 7.5 with 100 mM KCl.

For keratin, the inclusion bodies pellet was solubilized with 6 M Urea, 10 mM 2-Mercaptoethanol in 50 mM Tris HCl pH 7.5, clarified for 30 min at 10,000×*g*, at 15 °C, and the keratin-containing supernatant was collected. A size-exclusion chromatography using Superdex 200 Increase 10/300 GL (Cytiva) was performed and protein purity was checked using SDS-PAGE. Equimolar mixes of type-I and type-II keratins were incubated for 30 min at 22 °C in the above buffer, and heterotypic complexes of type-I and type-II keratins were separated from the uncomplexed proteins using an anion-exchange chromatography (HiTrapQHP, 1 mL, Cytiva) with a 25-mL gradient of 0-200 mM GuHCl in above buffer. Keratin filaments were then reconstituted in vitro by successive dialysis in 6 M Urea, 10 mM 2-Mercaptoethanol in 25 mM Tris HCl pH 7.5 for 4 hours, 2 M Urea, 5 mM 2-Mercaptoethanol in 5 mM Tris HCl pH 7.5 for 2 hours, and 5 mM 2-Mercaptoethanol in 5 mM Tris HCl pH 7.5 overnight. ^107^

### Fluorescent labeling of recombinant intermediate filament proteins

To produce fluorescently labelled keratin filaments, K14 was coupled to an Alexa fluor 488 C5-maleimide dye (Thermo Fisher scientific), complexed with a type-II keratin, and keratin filaments were reconstituted as described above. Fluorescently labelled vimentin was produced by coupling a Cy3b-maleimide dye (Cytiva) on the C-terminal cysteine of vimentin^C328A^-GGC prior to renaturation as described above.

Briefly, an inclusion bodies pellet of K14 or vimentin was solubilized with 6 M GuHCl, 50 mM Phosphate buffer pH 7.0, and clarified for 30 min at 10,000×*g*, at 4 °C. The protein-containing supernatant was purified using a size-exclusion chromatography using Superdex 200 Increase 10/300 GL (Cytiva) in the same buffer. Purity was checked by SDS-PAGE. Pure protein fractions were labelled with either Alexa fluor 488 C5-maleimide dye or Cy3b-maleimide dye, and subsequently the free dye was removed using a size-exclusion chromatography using Superdex 200 Increase 10/300 GL (Cytiva) in 6 M Urea, 10 mM 2-Mercaptoethanol in 50 mM Tris HCl pH 7.5 buffer for K14-Alexa488, or in 6M GuHCl in 10 mM TrisHCl pH 7.5 for vimentin-Cy3b. For K14, Alexa488-labelled heterotypic complexes were produced by mixing the Alexa488-labelled K14 with either unlabeled K5, or K6A, or K6Ah5rt, as described above. To produce Alexa488-labelled keratin filaments, unlabeled heterotypic complexes were mixed with 18% of the Alexa488-labelled ones, and filaments were then reconstituted in vitro by successive dialysis, as described above. For vimentin, the unlabeled wild-type vimentin was mixed with 37% of vimentin^C328A^-GGC-Cy3b. Note that the C328A substitution does not affect vimentin filament assembly. ^108^ The protein mix was then renatured as described above.

To determine the ability of labelled vimentin or keratin to form filaments, filamentous samples were inspected by negative stain electron microscopy. 3 µl of the filament solution was applied onto glow-discharged carbon-coated EM grids (G2300, Plano GmbH), incubated for 1 minute, and fixed with 1% uranyl acetate. Excess stain was removed by blotting. Specimen were examined in an FEI Tecnai G2 Spirit transmission electron microscope with a voltage of 120 kV.

### In-vitro co-sedimentation of recombinant intermediate filament proteins

Keratin filaments were reconstituted in using purified heterotypic complexes of K5/K14, K6A/K14, or K6A^h^5^rt^/K14, 18% labelled with Alexa Fluor 488 dye as described above. Unpolymerized, renatured vimentin, 37% labelled with Cy3b dye, was prepared separately. The three types of keratin filaments were incubated alone or with the unpolymerized vimentin for 30 min at 22 °C, and subsequently centrifuged at 20000×*g* for 30 minutes. Pellets were analyzed on a 9% SDS-PAGE and quantified by densitometry.

Alexa Fluor 488 and Cy3b fluorescence signals were imaged before Coomassie staining using FUSION FX7 Spectra (Vilber Smart Imaging) and quantified with Fiji software. Band fluorescence intensity measurements were made by integrating pixel densities in the band of interest and subtracting the contribution of the background. To quantify the ratio of vimentin co-sedimenting with keratin filaments, the band intensity of vimentin sedimented while incubated alone was subtracted from the total band intensity of vimentin sedimented in the presence of K5/K14, or K6A/K14, or K6A^h^5^rt^/K14 filaments, and subsequently divided by total band intensity of keratin in the corresponding pellet. Results were collected from two distinct experiments using the same batches of heterotypic complexes K5/K14, or K6A/K14, or K6A^h^5^rt^/K14, 18% labelled with Alexa Fluor 488 dye, and vimentin 37% labelled with Cy3b dye stored in urea or GuHCl, respectively.

### Electron microscopy of keratinocytes

For transmission electron microscopy of keratinocyte monolayers en face, glow-discharged carbon-film gold EM grids (EMS CF200-Au) were coated with bovine type I collagen (PureCol, Advanced Biomatrix #5005) by adding one drop of a 100 μg/mL collagen solution to each grid and incubating at 37 °C for 30 minutes. The collagen solution was then removed by absorption onto a strip of filter paper. Keratinocyte cell suspensions were prepared with 10^4^ cells/mL in differentiation medium, and one drop of cell suspension was added to each grid. After 24 hours, the grids were processed as follows^109^: grids were washed by dipping into 4 droplets of cytoskeleton buffer (CB; 10 mM MES, pH 6.1 with 150 mM NaCl, 5 mM EGTA, 5 mM glucose, and 5 mM MgCl_2_) ^110^, after wicking off the last droplet, the grids were lightly fixed and extracted in CB containing 0.25% glutaraldehyde and 0.5% Triton-X 100 for 1 minute. Grids were washed by dipping into 1 droplet of CB. The grids were then fixed for 10 minutes in CB containing 2% glutaraldehyde. After washing in 3 water droplets for 2 minutes each, the grids were stained with 1% uranyl acetate for 2 minutes; washed twice in water droplets for 2 minutes each, then air dried. Imaging was done on a JEM-1400 Plus transmission electron microscope equipped with a LaB6 source operated at 120 kV using an AMT-BioSprint 16M CCD camera. Cell junction widths were measured using ImageJ by manually drawing line regions of interest across the thickest portion of electron-dense intercellular bridges parallel to the cell-cell border.

For transmission electron microscopy of thin transverse sections of keratinocyte monolayers, cells were cultured in 35mm glass-bottom dishes (Mattek P35G-1.5-14-C) until confluent. The cells were washed once with PBS and twice with 100 mM sodium cacodylate buffer, then fixed in 2.5% glutaraldehyde in 100 mM cacodylate buffer. Following fixation, samples were rinsed five times in 100 mM sodium cacodylate, then post-fixed in 1% osmium tetroxide with 1.5% K_3_[Fe(CN)_6_] in 100 mM sodium cacodylate for 1 h at room temperature. Cells were rinsed with water and stained en bloc with 0.5% aqueous uranyl acetate in 25% methanol overnight at 4 °C. After five rinses with water, specimens were stained with 0.02 M lead nitrate in 0.03 M L-aspartate for 30 minutes. Samples were dehydrated with increasing concentration of ethanol, infiltrated with Hard Embed-812 resin and polymerized at 70 °C for 48 hours. Embed-812 discs were removed from the plastic housing by submerging the dish in liquid nitrogen. Paired disks of the same sample were then assembled with a drop of Hard Embed 812 resin at the center and polymerized for 24 hours. Blocks were sectioned with a diamond knife (Diatome) on a Leica Ultracut UCT(7) ultramicrotome (Leica Microsystems) and collected onto copper grids, post stained with 2% uranyl acetate in water and lead citrate. Images were acquired on a JEOL JEM-1400 Plus transmission electron microscope equipped with a LaB6 source operated at 120 kV using an AMT-BioSprint 16M CCD camera.

### Live imaging of monolayer migration

Sorted or mixed populations of keratinocytes were prepared depending on the experiment and seeded into collagen-coated 6-well tissue culture plates. Cells were grown to confluence, then the culture medium was changed from K-SFM to differentiation medium. After 24 hours, a linear scratch was created in each monolayer using a 1000 μL pipette tip. Monolayers were washed three times with PBS, which was then replaced with fresh differentiation medium. Samples were imaged using an inverted phase-contrast and epifluorescence Nikon ECLIPSE Ti microscope equipped with a motorized and programmable stage, a Nikon Perfect Focus System, a Hamamatsu Orca-Flash4.0 scientific CMOS camera, a SOLA solid state white-light excitation system, an OKO lab custom-built environmental chamber with temperature control and CO2 stage incubator, and Nikon Elements acquisition software. The environment was maintained at 37 °C with 5% CO_2_ during the imaging period. Beginning 30 minutes after wounding, phase-contrast and fluorescence images were acquired every 6 minutes using a Plan Fluor 10× 0.3NA objective.

Analysis of migration images was based on a previously established computer vision pipeline for migration velocity estimation and wound edge detection^38^. In brief, velocity measurements were computed from the phase-contrast images using a particle image velocimetry (PIV)-based method with a 15-μm square patch size. The pipeline includes a correction for possible microscope stage drift.

Monolayer contours were then determined by segmentation of the velocity map, based on the intuition that while the cells are moving, there should be no motion in the empty wound area. The algorithm imposes two additional constraints to increase robustness of the segmentation. First, the monolayer is assumed to be a continuous region. Second, the monolayer contour is assumed to move monotonically in one direction. The second condition means that the pipeline will fail if the monolayer edge retracts.

Monolayers where this occurred were excluded from the analysis. While the pipeline is robust to the presence of stationary debris or irregularities in the surface of the wound area, floating debris may interfere with velocity estimation and identification of the monolayer contour. Therefore, images with floating debris were also excluded from the analysis.

Downstream analysis steps were performed using custom MATLAB scripts. For uniform cell populations, two measures of migration were calculated. First, local migration speed, as estimated by particle image velocimetry, was averaged over an area between 10-μm and 200-μm from the monolayer contour. Because the monolayer contour as estimated from 15-μm PIV patches is necessarily imprecise, the 10-μm closest to the monolayer contour was excluded. Average local migration speed followed a consistent pattern, quickly increasing to a peak speed, then gradually declining over the course of the experiment (Figures S2B and 3C). However, the timing of this peak varied between individual scratch wounds. For more robust comparisons, migration speeds traces were aligned in time based on the time of peak migration speed for each wound. Second, area closed was calculated based on advancement of the monolayer contour. This measure was not adjusted for differences in timing of peak migration speed.

For mixed cell populations, monolayers were additionally segmented into regions with different keratin expression patterns. To accomplish this, each of the fluorescence images corresponding to tagged keratins were segmented using the Rosin thresholding method^111^. These masks were then used to define high and low expression regions for each keratin. Average local migration speeds were calculated separately for each keratin expression region in the 10-μm to 200-μm band from the monolayer contour, as well as for the band as a whole. As with the analysis for uniform cell populations, migration speed traces were aligned in time based on the time of peak migration speed. Since area closed is an overall measure that does not allow comparisons between different keratin expression regions, it was not considered in analysis of the mixed population monolayers.

### Fixed imaging of monolayer migration

Sorted or mixed populations of keratinocytes were prepared and seeded onto collagen-coated 18 mm-diameter #1.5 coverslips in tissue culture dishes. After cells were grown to confluence, the culture medium was switched from K-SFM to differentiation medium. After 4 hours, a line scratch was created in each coverslip using a 1000 μL pipette tip. The cultures were then returned to the incubator for 24 hours to allow migration to occur. After that period, coverslips were removed from the incubator, washed 3 times with PBS, incubated in 4% paraformaldehyde (Electron Microscopy Sciences #15713) in PBS for 10 minutes, incubated in 0.1% Triton X-100 in PBS for 8 minutes, and washed 3 more times with PBS. For experiments involving immunofluorescence labeling, samples were incubated in primary antibody diluted in PBS for 30 minutes at 37 °C (Table S2), washed 3 times with PBS, incubated in fluorophore-conjugated secondary antibody diluted in PBS for 30 minutes at 37 °C (Table S2), and washed 3 more times with PBS. Finally, coverslips were mounted onto glass slides using Fluoromount-G with DAPI (ThermoFisher 00-4959-52).

For analysis of relative enrichment of cells from a mixed population with different keratin expression levels at the monolayer edge (Figure S2E), slides were imaged using an inverted epifluorescence Nikon ECLIPSE Ti microscope equipped with a Zyla sCMOS camera (Andor), SOLA solid state white-light excitation system, and μManager acquisition software^112^. Partially overlapping images were acquired using a Plan Fluor 40× 1.3NA objective, then stitched together pairwise using the Fiji Image Stitching Plugin^113^. Analysis was performed on the composite stitched images. Monolayer edges were manually segmented using ImageJ. Individual cells were segmented using Cellpose version 1.0 with the pretrained “cyto” model^114^. A normalized sum of the tagged keratin fluorescence signals was used as input for the cytoplasm channel, and DAPI signal was used as input for the nuclear channel. Segmentations were manually reviewed, and any image areas with grossly incorrect cell segmentations were excluded from analysis. A custom Python script was used for downstream analysis. The median tagged keratin fluorescence signal was calculated for each cell. Cells were annotated as expressing high or low levels of each keratin using a linear discriminator function developed based on thresholds set between bimodal peaks in the fluorescence signal distributions. The fraction of cells expressing a high level of each keratin was then compared between the first 100-μm from the monolayer edge and the next 400-μm.

For immunofluorescence imaging of endogenous keratin expression in migrating monolayers (Figure 1D-E), slides were imaged using an inverted epifluorescence Nikon ECLIPSE Ti microscope equipped with a Zyla sCMOS camera (Andor), SOLA solid state white-light excitation system, and μManager acquisition software^112^. Images were acquired using Plan Apo 20× 0.75NA and Plan Apo 40× 1.3NA objectives. For immunofluorescence imaging of vimentin expression (Figure S6), slides were imaged using an inverted phase-contrast and epifluorescence Nikon ECLIPSE Ti microscope equipped with a motorized and programmable stage, a Nikon Perfect Focus System, a Hamamatsu Orca-Flash4.0 scientific CMOS camera, a SOLA solid state white-light excitation system, an OKO lab custom-built environmental chamber with temperature control and CO2 stage incubator, and Nikon Elements acquisition software. Images were acquired using a Plan Apo 60× 1.4NA objective.

### Live imaging of single-cell migration

Keratinocytes were sparsely seeded onto collagen-coated #1.5 glass-bottom culture dishes (Cellvis D35-20-1.5-N). Cells were imaged using an inverted phase-contrast and epifluorescence Nikon ECLIPSE Ti microscope equipped with a motorized and programmable stage, a Nikon Perfect Focus System, a Hamamatsu Orca-Flash4.0 scientific CMOS camera, a SOLA solid state white-light excitation system, an OKO lab custom-built environmental chamber with temperature control and CO2 stage incubator, and Nikon Elements acquisition software. The environment was maintained at 37 °C with 5% CO_2_ during the imaging period. Beginning 22-minutes, 35-minutes, or 6-hours after seeding, fluorescence images were acquired every 3 minutes for 6 to 18-hours using a Plan Fluor 10× 0.3NA objective.

Images were analyzed using the TrackMate plugin for Fiji^115^. Cells were identified using the Laplacian of Gaussian detector, and tracks were identified using the sparse linear assignment problem tracker^116^. Downstream analysis was performed using custom R scripts. Two complementary measures of migration speed were considered. First, mean velocity of a track was defined as the length of the track divided by the track duration. Second, mean squared displacement (MSD) rate was defined as the square of track displacement (linear distance from start to end positions) divided by the track duration, averaged over the set of tracks. For freely diffusing particles, MSD increases linearly with time at a rate proportional to the diffusion coefficient^117^. Since sparsely seeded cells migrating in the absence of a directional que do not fully resemble freely diffusing particles, we do not estimate a diffusion coefficient, and instead use the MSD rate as an approximation to aggregate information from multiple tracks of potentially different lengths.

### Stratified epidermal migration model

We adapted an epidermal culture model from existing protocols^118,119^. While epidermal equivalent cultures lack the fibroblast-containing dermal equivalent present in skin equivalent cultures ^96^, their simplified design allowed us to prepare them in removable barrier molds to model epidermal migration. To prepare epidermal cultures, polycarbonate filters with 0.4μm pore size (3.14cm^2^ area, Nunc #140640) were placed in larger tissue culture dishes. Two silicone inserts, each with two chambers separated by a 500μm gap (Ibidi #80209), were then pressed firmly into place on each filter using sterile forceps.

On day 0, a cell suspension of 2.5×10^5^ keratinocytes in 70μL of K-SFM was added into each chamber, and additional K-SFM was added to the culture dish to reach the level of the filter. Throughout the procedure, medium in the culture dish was refreshed every two to three days, and we verified that culture medium did not leak to the top surface of the filter outside of the silicone inserts. On day 3, culture medium was aspirated from above the filter within the silicone inserts, effectively lifting the epidermal culture to an air-liquid interface. At the same time, medium in the culture dish was changed from K-SFM to differentiation medium. On day 7, an additional 0.5 mM calcium chloride was added to the differentiation medium in the culture dish.

For migration experiments, the silicone inserts were removed on day 9 by grasping the insert and filter housing with sterile forceps and gently lifting the insert while holding the filter in place. The filter was then returned to the culture dish, maintaining epidermal cultures at an air-liquid interface during the migration period. This time point, 6 days after exposure of the culture to an air-liquid interface, corresponded with development of clear basal and suprabasal keratinocyte layers and an early cornified layer, but precedes the formation of a thick cornified layer, which was more easily disrupted during removal of the silicone inserts (Figure 1G). Cultures were either imaged live during migration or fixed at the appropriate timepoint.

### Fixation, staining, and imaging of epidermal cultures

Prior to fixation, medium was aspirated from the tissue culture dish and the top and bottom surfaces of the filters were washed twice with PBS, taking care not to dislodge the epidermal cultures from the top surface of the filters. The filters were then removed from the culture dishes and incubated overnight in 4% paraformaldehyde solution at 4 °C (Electron Microscopy Sciences #15713). In preparation for paraffin embedding and sectioning, the filters were cut out from their plastic housing using a #15 scalpel blade (GF Health Products 2976#15), bisected perpendicularly to the migration edge, and embedded in a 2.5% agarose gel (Lonza SeaKem LE Agarose #50000). Samples were submitted to the UT Southwestern Histo Pathology Core Facility for paraffin embedding, sectioning, and preparation of hematoxylin and eosin (H&E) stained and unstained slides.

In preparation for immunofluorescence staining, unstained paraffin-embedded tissue sections on glass slides were deparaffinized by 3 successive 5-minute incubations in xylene followed by rehydration in 3 successive 3-minute incubations in 100% ethanol and 2 successive 2-minute incubations in deionized water. For antigen retrieval, slides were placed in Tris-EDTA buffer (10 mM Tris base, 1 mM EDTA, 0.05% Tween 20, pH 9), heated to 95 °C in a microwave oven, and incubated until the buffer cooled to room temperature, approximately 45 minutes. Slides were rinsed three times in Hanks Balanced Salt Solution (HBSS) for 5 minutes each, and a PAP pen was used to circle the tissue sections. Slides were then incubated in blocking solution [5% goat serum (Abcam ab7481), 5% bovine serum albumin (Equitech-Bio BAH65-0500), and 0.5% Triton X-100 (Sigma X100) in PBS] for 1 hour. Primary antibodies (Table S2) were diluted in PBS and incubated on the tissue sections overnight at 4 °C in a humidified chamber. Following primary antibody incubation, slides were washed 3 times with HBSS for 10 minutes each. Secondary antibodies (Table S2) were diluted in PBS and incubated on the tissue sections for 60 minutes. Finally, the slides were washed 3 times with HBSS for 10 minutes each, and a small amount of Fluoromount-G with DAPI (ThermoFisher 00-4959-52) was used to affix coverslips to the slides.

H&E slides were imaged using an Olympus BX53 upright microscope with brightfield illumination, an Olympus DP21 2.01 megapixel color CCD camera, and Plan N 4× 0.1NA, 10× 0.25NA, 20× 0.4NA, and 40× 0.65NA objectives. For migration assays, the gap width between epidermal culture pairs was measured using ImageJ by manually drawing polyline regions of interest along the filter in the gap.

Immunofluorescence slides were imaged using an inverted epifluorescence Nikon ECLIPSE Ti microscope equipped with a Zyla sCMOS camera (Andor), SOLA solid state white-light excitation system, and μManager acquisition software^112^. Partially overlapping images were acquired using a Plan Fluor 40× 1.3NA objective, then stitched together pairwise using the Fiji Image Stitching Plugin^113^.

### Live imaging of epidermal culture migration

Epidermal cultures were imaged using an inverted phase-contrast and epifluorescence Nikon ECLIPSE Ti microscope equipped with a motorized and programmable stage, a Nikon Perfect Focus System, a Hamamatsu Orca-Flash4.0 scientific CMOS camera, a SOLA solid state white-light excitation system, an OKO lab custom-built environmental chamber with temperature control and CO2 stage incubator, and Nikon Elements acquisition software. With epidermal cultures on filters sitting in tissue culture dishes placed on the stage of the inverted microscope, images were acquired from below, through the semi-translucent filter (Figure 2H). The environment was maintained at 37 °C with 5% CO_2_ during the imaging period. Beginning 15 minutes after barrier removal, fluorescence images were acquired every 12 minutes using a Plan Fluor 10× 0.3NA objective.

Although resolution was limited by imaging through the filters, bright fluorescence signals from the tagged keratin constructs allowed us to identify cell locations. To analyze the resulting fluorescence images, we modified the monolayer migration analysis pipeline described above^38^ as follows: First, all images were smoothed using a Gaussian filter with a 3.25-μm (5 pixel) standard deviation to decrease the influence of the filter mesh pattern on the PIV calculations. Second, since the initial texture-based segmentation seed frequently failed with fluorescence images instead of phase-contrast images, we used Otsu thresholding as an alternative^120^. Finally, because interpretation of the PIV-based local velocity estimates was complicated by potential differences in cell motion at different vertical levels within the three-dimensional epidermal culture, migration distance and speed calculations were based only on the distance travelled by the monolayer contour.

### Keratin filament network segmentation and analysis

Keratinocyte cell lines expressing fluorescently tagged keratins were seeded onto collagen-coated #1.5 glass-bottom culture dishes (Cellvis D35-20-1.5-N). For knockdown experiments, cells were transduced with lentivirus to express targeting shRNA (Table S1) or a scrambled control 72 hours before analysis. Once cells grew to discrete colonies, but before reaching confluence, the culture medium was switched from K-SFM to differentiation medium. After 8 hours, cells were imaged using an inverted phase-contrast and epifluorescence Nikon ECLIPSE Ti microscope equipped with a motorized and programmable stage, a Nikon Perfect Focus System, a Hamamatsu Orca-Flash4.0 scientific CMOS camera, a SOLA solid state white-light excitation system, an OKO lab custom-built environmental chamber with temperature control and CO2 stage incubator, and Nikon Elements acquisition software. The environment was maintained at 37 °C with 5% CO_2_ during the imaging period. Images were acquired every 1 minute for 4 minutes using a Plan Apo 60× 1.4NA objective, generating an effective pixel size in object space of 108 nm. For applicable experiments, 30 μM Blebbistatin (Sigma B0560), 5 μM Y27632 (Selleckchem S1049), 1 μM GSK 269962A (Selleckchem S7687), or DMSO (Fisher BP231) control were added 1 hour (Blebbistatin) or 2 hours (other inhibitors) prior to imaging.

Images were analyzed using the previously published Filament Network Analysis package written in MATLAB^39^. In brief, the pipeline segments the filament network using a multi-scale steerable filter and iterative graph matching to connect fragments into complete filaments. The pipeline also calculates the local structural similarity of two filament networks based on the distance and orientation of individual filament pixels in the networks. Each pixel in the first network is matched to the closest pixel in the second network within a defined radius, and vice versa. A radius of 2.2 μm was used for all studies here. For each matched pair, the difference in position and filament orientation are calculated and used to create distance and angle maps, which are spatially smoothed by a Gaussian filter, combined, and scaled so the final similarity score decreases monotonically from 1 to 0 as the distance increases to the search radius or the angle increases to π/3. The dynamics score is defined as 1 minus the combined similarity score. The final dynamics score is 0 for identical networks and increases to 1 with increasing local differences in filament position or orientation.

To measure the effect of keratin filament composition on filament network dynamics within cells, the keratin filament images at baseline were compared to images 1, 2, 3, or 4 minutes later (Figure 4A-B). Because this analysis cannot reliably distinguish movement of filaments within cells from movement of the cells themselves, all image sequences were pre-screened to ensure that only stationary cells were included in the analysis. Cells were manually segmented using ImageJ, and the Filament Network Analysis Package was run to segment the keratin filament networks and calculate dynamics score maps (Figure 4C-D). The dynamics score map was then averaged over each cell area mask, excluding regions without a filament match. As a check, a quality score was computed for each cell, defined as the fraction of the cell mask area with a filament match, which was typically higher than 0.9. Image sequences of cells with low quality scores were reviewed by visual inspection. They tended to show failed filament segmentation due to low signal-to-noise ratio, debris, or cell movement, which was missed in the initial screen; these cells were excluded from subsequent analysis. For the filament dynamics analysis of keratinocytes expressing different levels of multiple keratins, the quality score on each channel also served as a robust marker of high versus low expression of each keratin, with clear bimodal distributions (Figure S3J).

In addition to the dynamics score, the Filament Network Analysis package was used to calculate additional keratin filament network properties from the baseline keratin filament segmentation. Filament density was defined as the fraction of cell area pixels segmented as filaments. Filament straightness was defined as the ratio of the linear start to end distance to the filament length. Filament curvature was defined as the mean curvature of a polynomial fit at each pixel along the filament. The median filament length, straightness, and curvature were calculated for each cell. An alternate cell summary measure of filament curvature was also calculated as the median curvature of each pixel in all filaments in the cell.

### Traction force microscopy

Silicone gel substrates for traction force microscopy (TFM) were prepared in #1.0 glass-bottom 35-mm tissue culture dishes (Cellvis D35-14-1.0-N). Prepolymers for high-refractive index (*n* = 1.49) 8-kPa gels were prepared by mixing components A and B of QGel 920 (Quantum Silicones) in a 1:1.15 ratio as previously described^43,121^. The prepolymers were spread onto the coverglass bottoms of the dishes by spinning at 1,250 r.p.m for 30 seconds in a spin-coater (Laurell Technologies WS-650Mz-23NPPB), then cured at 100 °C for 2 hours, producing gels approximately 35-μm thick. The gels were then treated with 5% (3-aminopropyl)triethoxysilane (APTES) in ethanol to functionalize their surfaces. To allow for visualization and tracking of substrate deformation, 40-nm carboxylate far-red fluorescent beads were covalently linked to the gel surfaces by incubating the gels under a suspension of the beads (1:5,000 dilution from the 5% stock suspension) in 40-mM HEPES, pH 8, with 0.01% 1-ethyl-3-(3-dimethylaminopropyl) carbodiimide (EDC) as a catalyst. To facilitate cell adhesion during cell seeding, the substrates were coated with fibronectin by incubation with 50-μg/ml of fibronectin and 1 mg/ml EDC in PBS for 15 minutes at room temperature. The coated dishes were washed three times with PBS and filled with cell culture medium before cell seeding.

To measure keratinocyte traction force generation, 4×10^4^ cells were seeded onto a substrate in differentiation medium, then returned to the incubator for 4.5 hours to allow attachment to occur. For knockdown experiments, cells were transduced with lentivirus to express targeting shRNA or a scrambled control 72 hours before seeding onto TFM substrates. Following attachment, cells were imaged using a GE DeltaVision OMX SR inverted microscope equipped with a motorized and programmable stage, three PCO sCMOS cameras, solid state lasers (405nm, 488nm, 568nm, and 640nm), and environment control. Images of the substrate beads and keratin tags were acquired in Ring-TIRF mode with a U Plan Apo 60× 1.49NA objective (generating an effective pixel size in object space of 80 nm), with imaging performed at 37 °C. Following initial imaging, during which cell positions were recorded, a prewarmed solution of dilute bleach was added to remove cells from the substrate, thereby allowing repeat imaging of the same locations with the substrate in its relaxed, strain-free state.

Image analysis was performed using the previously published TFM package written in MATLAB^43,122^. Briefly, the substrate deformation field was determined by tracking the displacement of each bead between the deformed and relaxed images using cross-correlation analysis. The displacement field, after outlier removal and stage drift correction, was then used to estimate the traction field over areas of interest surrounding each cell. Traction fields were estimated using the boundary element method^43^.

While more computationally intensive than the more commonly used Fourier transform traction cytometry method, the boundary element method allows L1-norm rather than L2-norm regularization, which resolves force variation at shorter length scales^43^. A regularization parameter of 0.05 was selected based on L-curve analysis of a subset of images^43^. This parameter value was applied to all images analyzed to permit comparison of the resulting force fields. Cell area was segmented based on the tagged keratin signal. Since this signal did not always clearly extend to the cell boundary (for example, Figure 5A), masks were dilated by 1.6 μm (20 pixels), although we verified that the precise amount of dilation did not meaningfully affect the results (Figure S4C). Strain energy, which represents the mechanical work of the cell on the elastic substrate, was calculated as ½ × (displacement · traction), integrated over the segmented cell area^123^.

### Proximity ligation assay

Mixed populations of keratinocytes were prepared and seeded onto 8-chamber glass slides with removable wells (Lab-Tek, ThermoFisher #154534). Cells were grown to approximately 50% confluence, then culture medium was switched from K-SFM to differentiation medium for 24 hours. For knockdown experiments, cells were transduced with lentivirus to express targeting shRNA (Table S1) or a scrambled control 72 hours before processing. Slides were removed from the incubator and washed three times with PBS. For experiments using antibodies against keratins, the slides were incubated in methanol for 2 minutes. For other experiments, the slides were incubated in 4% paraformaldehyde in PBS for 10 minutes, then incubated in 0.1% Triton X-100 in PBS for 8 minutes. Slides were then washed twice with PBS and processed using Duolink Proximity Ligation Assay (PLA) reagents (In-Situ PLA Probe Anti-Mouse MINUS, Sigma DUO92004; In-Situ PLA Probe Anti-Rabbit PLUS, Sigma DUO92002; In-Situ Detection Reagents FarRed, Sigma DUO92013) according to the manufacturer’s in situ fluorescence protocol. Briefly, slides were incubated in the provided Blocking Solution for 60 minutes, incubated in primary antibodies diluted in the provided Antibody Dilutant solution for 30 minutes (Table S2), incubated in PLA probe solution for 60 minutes, incubated in the ligation solution for 30 minutes, and incubated in the amplification solution for 100 minutes, with washes in the provided buffers between each step. Finally, a #1.5 cover glass was affixed to the slide using Duolink In-Situ Mounting Medium with DAPI (Sigma DUO82040).

Slides were imaged using an inverted phase-contrast and epifluorescence Nikon ECLIPSE Ti microscope equipped with a motorized and programmable stage, a Nikon Perfect Focus System, a Hamamatsu Orca-Flash4.0 scientific CMOS camera, a SOLA solid state white-light excitation system, an OKO lab custom-built environmental chamber with temperature control and CO2 stage incubator, and Nikon Elements acquisition software. Images were acquired on the tagged keratin, PLA reaction, and DAPI channels using a Plan Apo 60× 1.4NA objective (generating an effective pixel size in object space of 110 nm).

Image analysis was performed using custom Python and MATLAB scripts. Individual cells were segmented using Cellpose version 1.0 with the pretrained “cyto” model^114^. A normalized sum of the tagged keratin fluorescence signals was used as input for the cytoplasm channel, and DAPI signal was used as input for the nuclear channel. The median tagged keratin fluorescence signal was calculated for each cell. Cells were annotated as expressing high or low levels of each keratin using a linear discriminator function developed based on thresholds set between bimodal peaks in the fluorescence signal distributions. Segmentations and classifications were manually reviewed, and any incorrectly segmented or classified cells were excluded from analysis. Cells that were poorly adhered to the slide or out of focus were also excluded. Next, individual local intensity clusters in the scaled PLA image were detected using previously published MATLAB code combining wavelet denoising and multiscale products of wavelet coefficients^124,125^. Finally, PLA detections were counted within each segmented cell mask, and the density of PLA detections was compared between cells with different keratin expression levels.

### Proximity biotin labeling and proteomics

Keratinocytes were transduced with lentivirus to express K5 or K6A linked to a small promiscuous biotin ligase^54^. Because these constructs lack a selection marker, a mixed-population lentivirus was prepared by combining the tagged keratin constructs with an EGFP marker construct (see “Plasmids,” above) at a molar ratio of 100:1. Cells were selected by fluorescence activated cell sorting as described above. Each cell line was then grown to confluence in collagen-coated 10 cm tissue culture plates and switched from K-SFM to differentiation medium for 24 hours before beginning the experiment.

Two independent experimental replicates were performed. Cells were incubated with 100 µM biotin for 1 hour and lysed in JS buffer (50 mM Tris HCl pH7.5, 150 mM NaCl, 5 mM EGTA, 1.5 mM MgCl_2_, 1% Glycerol and 1% Triton X-100) supplemented with protease and phosphatase inhibitors (Halt Protease and Phosphatase Inhibitor, Thermo). Cleared lysates were incubated with Protein A Sepharose beads to remove proteins with nonspecific binding to the beads. The resulting supernatant (1 mg) was incubated with 100 µl of streptavidin-coated magnetic beads (Dynabeads MyOne Streptavidin T1 beads, Invitrogen) overnight at 4 °C with rotation to enrich for biotinylated proteins. Beads were washed three times with ice-cold JS buffer and in-gel digested for Mass Spectrometry analysis.

Proteomics data was analyzed using the DEP package for differential enrichment analysis of proteomics data in R^126^. Only proteins with no missing values in at least one of the two cell lines were included in the analysis. Abundance data was background-corrected and normalized by variance stabilizing transformation^127^, and any remaining missing values were imputed as draws from a Gaussian distribution centered on quantile 0.005 of the observed values with standard deviation equal to the median standard deviation of the observed values^128^. Differential enrichment was estimated based on protein-wise linear models and empirical Bayes statistics^129^, and local false discovery rates were estimated using the fdrtool package^130,131^. Functional analysis was then performed using the clusterProfiler package^132^. We searched for overrepresentation of proteins with at least 3-fold enrichment and adjusted *P* < .05 in the Gene Ontology Biological Process annotation set^133,134^ using Fisher’s exact test, with adjustment for multiple comparisons^135^. Because the small size of the screen limited our statistical power, we restricted our search to high-level labels with between 750 and 5,000 annotated genes.

### Analysis of cell-cell junction density

Cells were prepared and seeded onto collagen-coated 18-mm-diameter #1.5 coverslips in tissue culture dishes. After reaching 25% confluence, the culture medium was switched from K-SFM to differentiation medium. After 5 hours, coverslips were removed from the incubator, washed 3 times with PBS, incubated in 4% paraformaldehyde (Electron Microscopy Sciences #15713) in PBS for 10 minutes, incubated in 0.1% Triton X-100 in PBS for 8 minutes, and washed 3 more times with PBS. Coverslips were then incubated in either Alexa Fluor 488 phalloidin (ThermoFisher A12379) or Alexa Fluor 555 phalloidin (ThermoFisher A34055) diluted 1:400 in PBS for 30 minutes at 37 °C. Finally, coverslips were mounted onto glass slides using Fluoromount-G with DAPI (ThermoFisher 00-4959-52). Slides were imaged using an inverted Nikon Ti-Eclipse microscope equipped with a pco.edge sCMOS camera with 6.5-μm pixel size (PCO), an Andor Diskovery illuminator coupled to a Yokogawa CSU-X1 confocal spinning disk head with 100 nm pinholes, and an Apo TIRF 60× 1.49NA objective (Nikon) with an additional 1.8× tube lens (yielding a final magnification of 108×; Andor Technology). This setup generated an effective pixel size in object space of 120 nm. Actin and keratin filament networks were segmented using the Filament Network Analysis package as described above in the section “Keratin filament network segmentation and analysis” based on images of phalloidin and tagged keratins respectively^39^. Approximate cell-cell boundaries were manually annotated using ImageJ. Adherens junctions and desmosomes, defined as segmented actin and keratin filaments crossing annotated cell-cell boundaries, were counted using the SlideSet plugin with a custom analysis plugin written in Java^136^. Junction density was defined as the number of junctions detected along a cell-cell boundary divided by the length of the boundary. To account for potential inconsistency in the start and end positions of the manually annotated cell-cell boundaries, boundary length was trimmed by the outermost junctions. Junction density was then compared between cell lines with different keratin expression levels.

### Cell death assay

Cells were prepared and 5.5×10^4^ cells were seeded onto collagen-coated 35-mm #1.5 glass-bottom culture dishes (Cellvis D35-20-1.5-N). After 4 hours of incubation, cells were transduced with lentivirus to express targeting shRNA or a scrambled control. Cells were cultured with shRNA expression for 120 hours with regular culture medium changes. H2B-mCherry3 co-expression from the shRNA constructs allowed us to confirm an infection efficiency close to 100% (see “Plasmids” above). Following the growth period, cells were incubated in SYTOX Deep Red Nucleic Acid Stain dissolved in K-SFM (0.5 μM final concentration; ThermoFisher S11381) for 30 minutes at 37 °C, then washed twice with PBS. Cells were imaged using an inverted epifluorescence Nikon ECLIPSE Ti microscope equipped with a motorized and programmable stage, a Nikon Perfect Focus System, a Hamamatsu Orca-Flash4.0 scientific CMOS camera, a SOLA solid state white-light excitation system, an OKO lab custom-built environmental chamber with temperature control and CO2 stage incubator, and Nikon Elements acquisition software. Images of the SYTOX dye, tagged keratins, and tagged H2B were acquired with a Plan Apo 60× 1.4NA objective, generating an effective pixel size in object space of 108 nm.

Individual cells were segmented using Cellpose version 1.0 with the pretrained “cyto” model^114^. First, the tagged keratin fluorescence signal was used as input for the cytoplasm channel, and no input was provided for the nuclear channel. Second, the tagged H2B signal was used as input for the “cytoplasm” channel, and no input was provided for the “nuclear” channel. This produced whole cell and nuclear masks respectively. Segmentations were manually reviewed, and any image areas with grossly incorrect cell segmentations were excluded from analysis. A custom Python script was used for downstream analysis. The median SYTOX signal within the nuclear mask was calculated for each whole cell mask.

Cells were annotated as either SYTOX-negative (live) or SYTOX-positive (dead) based on comparison of the median SYTOX signal to a threshold set between bimodal peaks in the SYTOX signal distribution.

### Cell proliferation assay

Cells were prepared and seeded into collagen-coated 12-well tissue culture plates, 4×10^4^ cells per well. For knockdown experiments, cells were transduced with lentivirus to express targeting shRNA or a scrambled control after 4 hours, then analyzed after 96 hours with regular changes of culture medium. H2B-mCherry3 co-expression from the shRNA constructs allowed us to confirm an infection efficiency close to 100% (see “Plasmids” above). For direct cell line comparisons without shRNA knockdown, cells were analyzed after 4 days in standard culture conditions. Following the growth period, cells were incubated in 5-ethynyl-2’-deoxyuridine (EdU; ThermoFisher A10044) dissolved in K-SFM (20 mM final concentration) for 60 minutes at 37 °C. Cells were then washed 3 times with PBS, incubated in 4% paraformaldehyde (Electron Microscopy Sciences #15713) in PBS for 10 minutes, incubated in 0.1% Triton X-100 in PBS for 8 minutes, and washed 3 more times with PBS. To detect EdU incorporation into nuclear DNA, cells were incubated in a reaction mixture containing 20 mg/mL L-ascorbic acid (Sigma A92902), 0.5 mg/mL anhydrous copper(II) sulfate (Acros #42287), and 1 μM Cy5-azide (Sigma #777323) in PBS for 30 minutes at 37 °C. Cells were washed 3 times with PBS, then incubated in DAPI (ThermoFisher D1306) diluted 1:5,000 in PBS for 15 minutes at room temperature to provide a nuclear counter-stain. After 3 final washes in PBS, cells were imaged using an inverted phase-contrast and epifluorescence Nikon ECLIPSE Ti microscope equipped with a motorized and programmable stage, a Nikon Perfect Focus System, a Hamamatsu Orca-Flash4.0 scientific CMOS camera, a SOLA solid state white-light excitation system, an OKO lab custom-built environmental chamber with temperature control and CO2 stage incubator, and Nikon Elements acquisition software. DAPI and Cy5 images were acquired using a Plan Fluor 10× 0.3NA objective, generating an effective pixel size in object space of 650 nm.

Images were segmented using Cellpose version 1.0 with the pretrained “cyto” model^114^. The DAPI signal was used as input for the “cytoplasm” channel, and no input was provided for the “nuclear” channel, producing cell nucleus masks. The median Cy5 signal was calculated for each mask, and cells were annotated as EdU-positive (Cy5-high, proliferating) or EdU-negative (Cy5-low, quiescent) based on comparison of the median Cy5 signal to a threshold set between bimodal peaks in the Cy5 signal distribution.

### Imaging of clinical wound samples

Routine skin biopsies performed for diagnostic purposes result in epidermal wounds, which typically heal over 2 to 6 weeks. If a malignant lesion is identified in the biopsy, an excision may be recommended to ensure that the malignancy is completely removed. If an excision is performed before the biopsy wound has completely healed, the resulting specimen will incidentally contain the re-epithelializing biopsy wound edges. We obtained anonymized H&E stained and unstained tissue sections from an excision specimen in which the re-epithelizing biopsy wound edges were visible, but no residual carcinoma was present. Unstained slides were processed for immunofluorescence and imaged using the procedures describe above in “Fixation, staining, and imaging of epidermal cultures.” This study was approved by the UT Southwestern Institutional Review Board.

### Clinical sample transcriptomics analysis

Two previously published transcriptomics datasets comparing wounded to intact skin were reanalyzed for keratin expression using custom R scripts. Iglesias-Bartolome and colleagues collected serial punch biopsy specimens from the arm and buccal mucosa, extracted RNA from the samples, and performed RNA sequencing^24^. Unwounded skin was represented by the initial samples, and wounded skin was represented by follow-up samples taken from the same locations. We downloaded the expression data (Gene Expression Omnibus GSE97615) for the arm skin samples and plotted keratin transcript levels (log-transformed RPKM). Statistical analysis was not performed. Ramirez and colleagues collected tissue samples from diabetic foot ulcers and non-ulcerated foot skin, extracted RNA from the samples, and profiled transcripts using Affymetrix GeneChip Human Gene 2.0 ST microarrays^25,137^. We downloaded the expression data (Gene Expression Omnibus GSE80178) and used the “oligo” R package to import, annotate, and preprocess the microarray data, including background-correction and normalization using the RMA algorithm^138^. Keratin expression intensity values for wound and control samples were plotted, and statistical analysis was not performed.

### Statistical analysis

All statistical analysis was performed using R (R Foundation for Statistical Computing). The Kruskal-Wallis rank sum test was used to evaluate non-parametric scaled data, with Dunn’s method for multiple comparisons used for experiments involving more than two groups^139^. Two-tailed *P* values less than 0.05 are considered to be significant and are shown in figures or indicated in legends.

## Supplementary Information

**Table S1.**
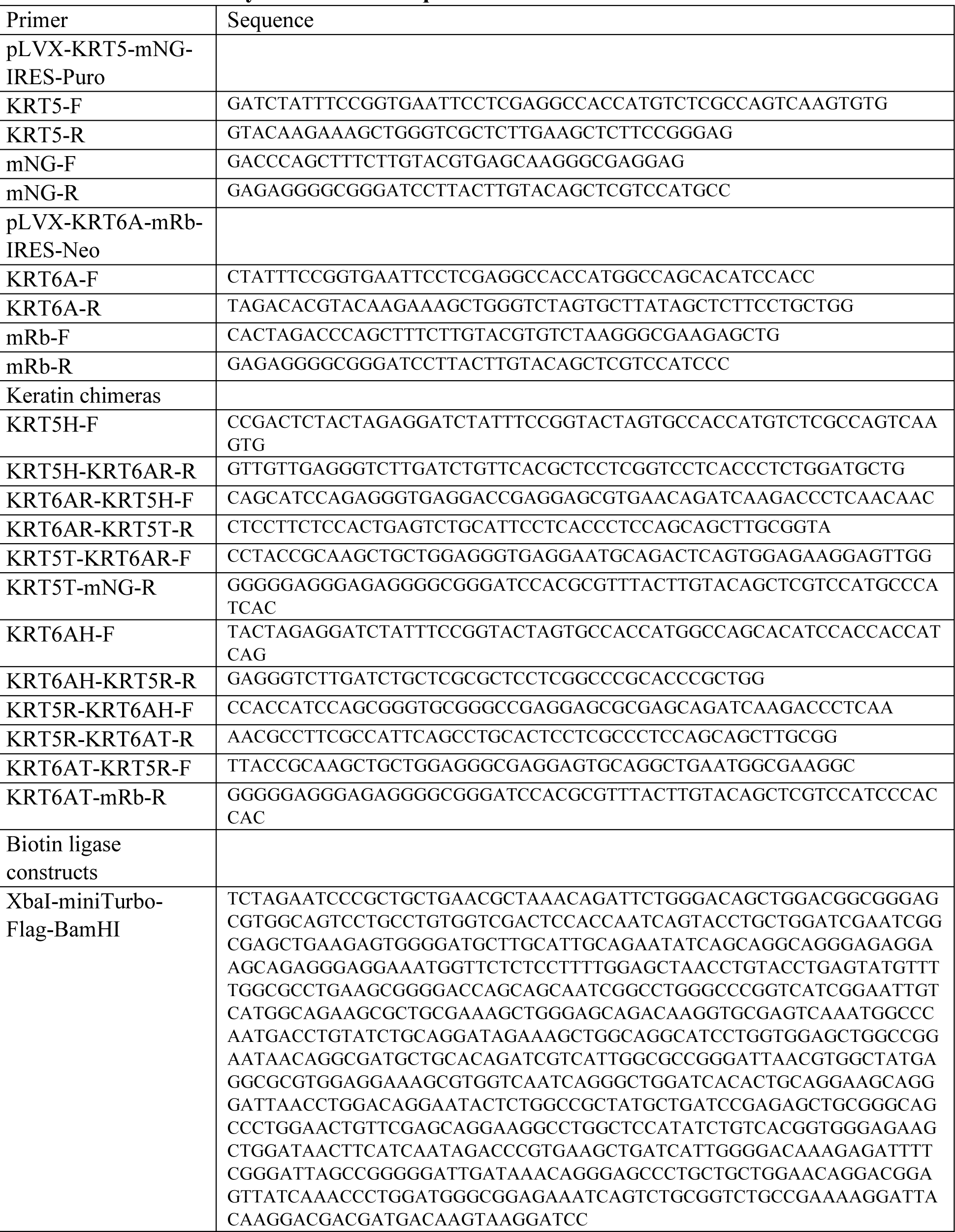
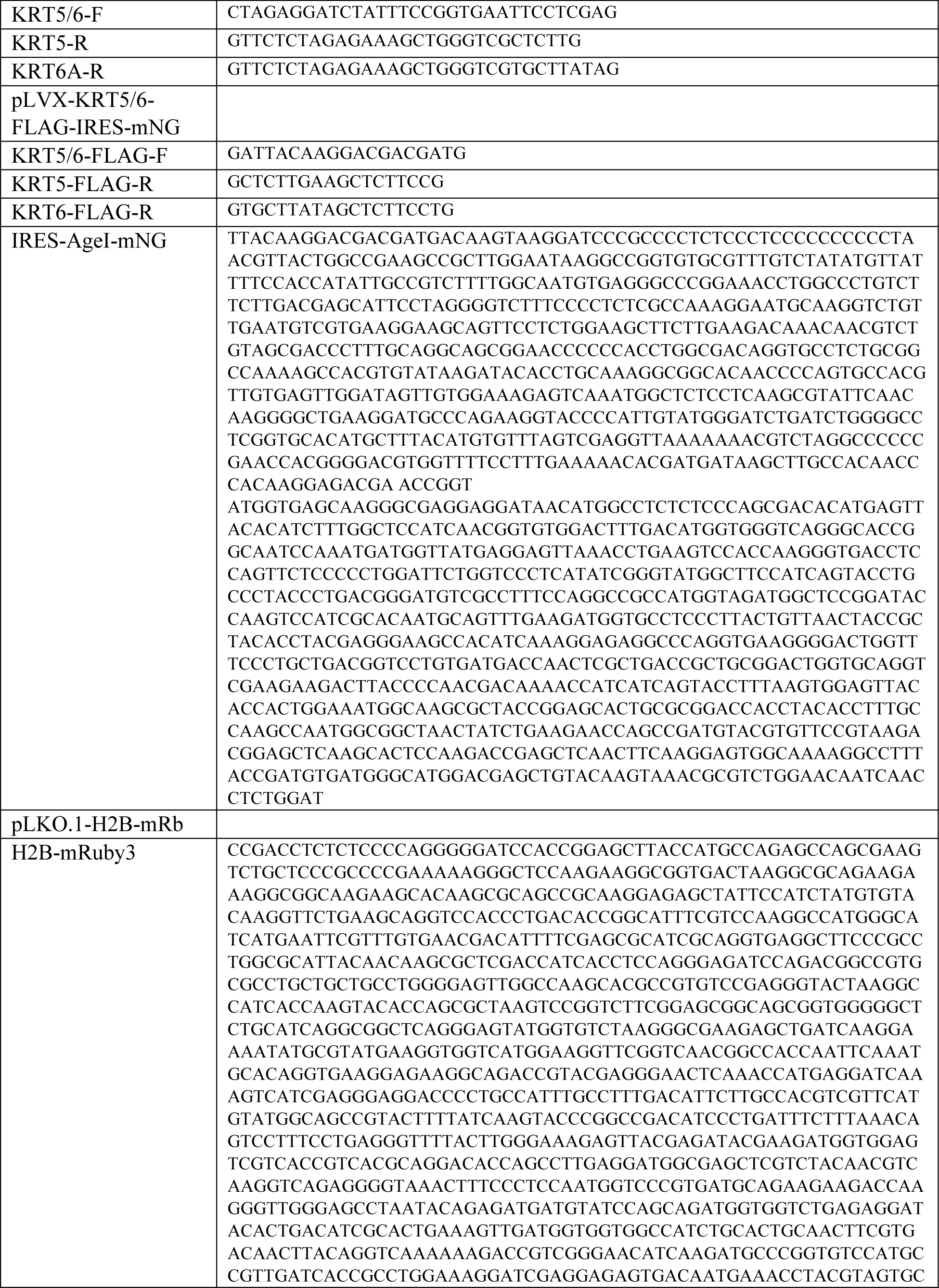
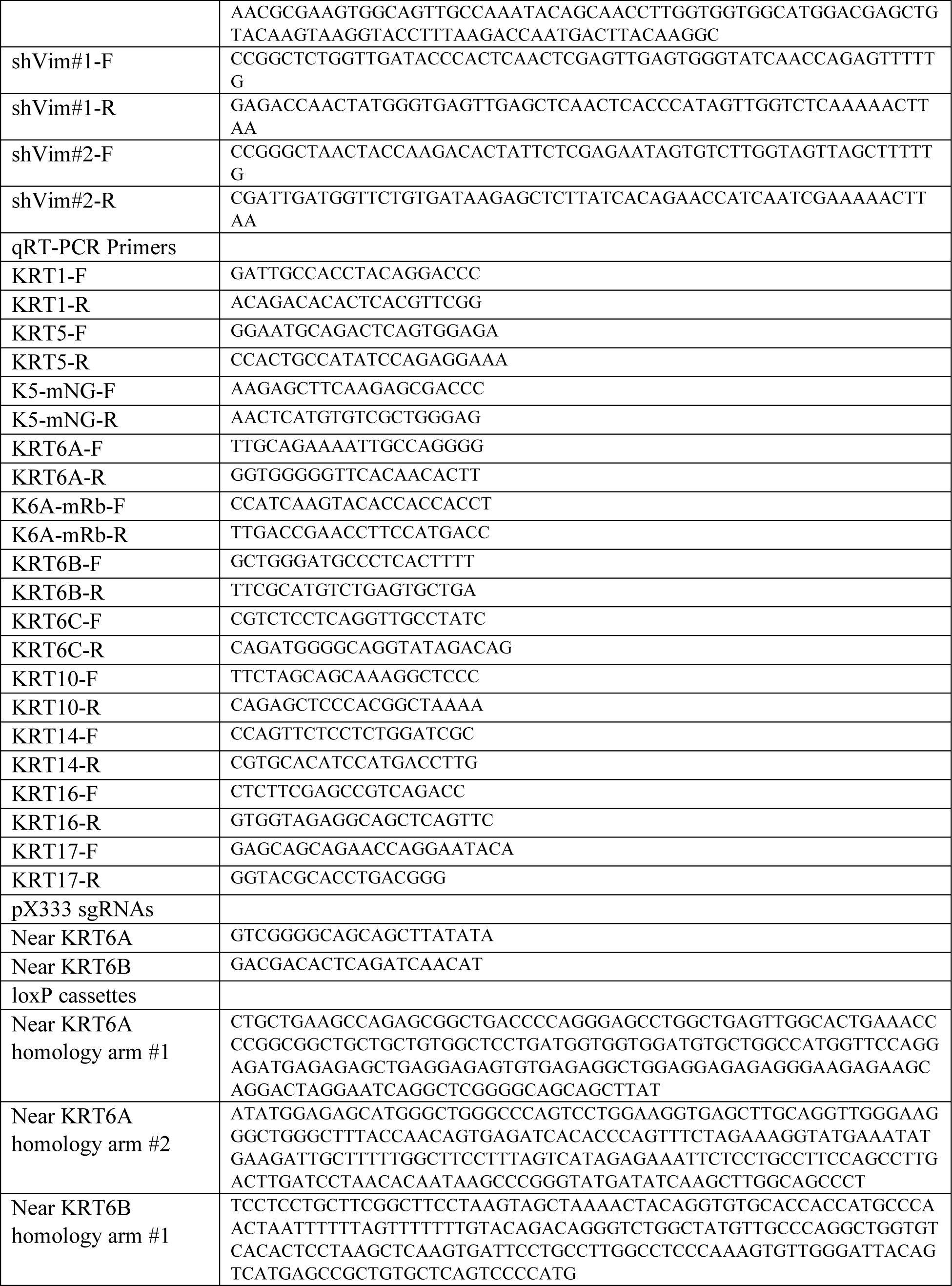
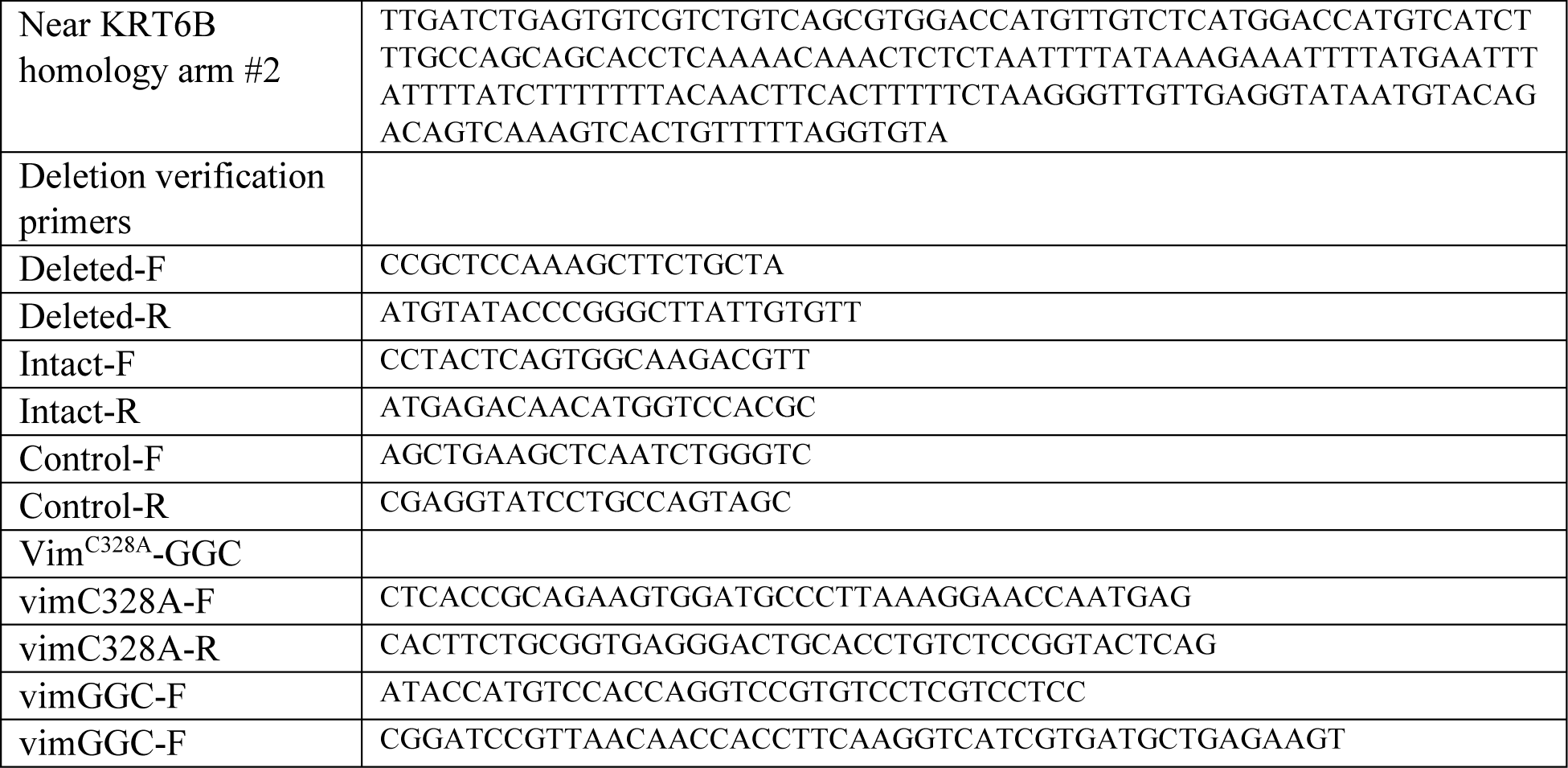
Primers and synthetic DNA sequences

**Table S2.** Antibodies

| Antibody | Application and dilution |
| --- | --- |
| Mouse anti-keratin 1 (Novus LHK1) | WB, 1:250 |
| Rabbit anti-keratin 5 (Abcam ab52635), N-terminal epitope | IF/PLA, 1:200<br>WB, 1:10,000<br>IHC, 1:200 |
| Rabbit anti-keratin 5 (Cell Signaling Technology #25807), C-terminal epitope | WB, 1:1,000 |
| Rabbit anti-keratin 6 (Abcam ab93279), epitope conserved in K6A, K6B, and K6C | IF, 1:50<br>WB, 1:2,000<br>IHC, 1:50 |
| Rabbit anti-keratin 10 (Abcam ab76318) | WB, 1:1,000 |
| Mouse anti-keratin 14 (EMD Millipore MAB3232) | WB, 1:200 |
| Mouse anti-keratin 16 (Invitrogen LL025) | WB, 1:500 |
| Rabbit polyclonal anti-keratin 17 (Abcam ab53707) | IF/PLA, 1:1,000<br>WB, 1:1,000<br>IHC, 1:100 |
| Mouse anti-beta-actin (Sigma A1978) | WB, 1:5,000 |
| Rabbit polyclonal anti-non-muscle myosin heavy chain IIA (MYH9; Invitrogen PA5-17025) | PLA, 1:200 |
| Mouse anti-phospho-myosin regulatory light chain (Cell Signaling Technology #3675) | PLA, 1:100 |
| Mouse anti-vimentin (Sigma V6630) | IF/PLA, 1:100 |
| Rabbit anti-vimentin (Cell Signaling Technology #5741) | WB, 1:1,000 |
| Rabbit anti-ROCK2 and ROCK1 (Abcam ab45171) | PLA, 1:250 |
| Rabbit polyclonal anti-myosin light chain kinase (Invitrogen PA5-76982) | PLA, 1:200 |
| HRP, goat anti- mouse IgG (Invitrogen #31430) | WB, 1:1,000 |
| HRP, goat anti- rabbit IgG (Invitrogen #31460) | WB, 1:1,000 |
| Alexa Fluor 488, goat anti- mouse IgG (Invitrogen A-11209) | IF, 1:500 |
| Alexa Fluor 488, goat anti- rabbit IgG (Invitrogen A-11008) | IF, 1:500 |
| Alexa Fluor 568, goat anti- mouse IgG (Invitrogen A-11004) | IF, 1:500 |
| Alexa Fluor 568, goat anti- rabbit IgG (Invitrogen A-11036) | IF, 1:500 |
| Alexa Fluor 647, goat anti- rabbit IgG (Invitrogen A-21244) | IF, 1:500 |

IF, immunofluorescence; IHC, immunohistochemistry or immunofluorescence on formalin-fixed paraffin-embedded samples; PLA, proximity ligation assay; WB, western blot

**Figure S1.**
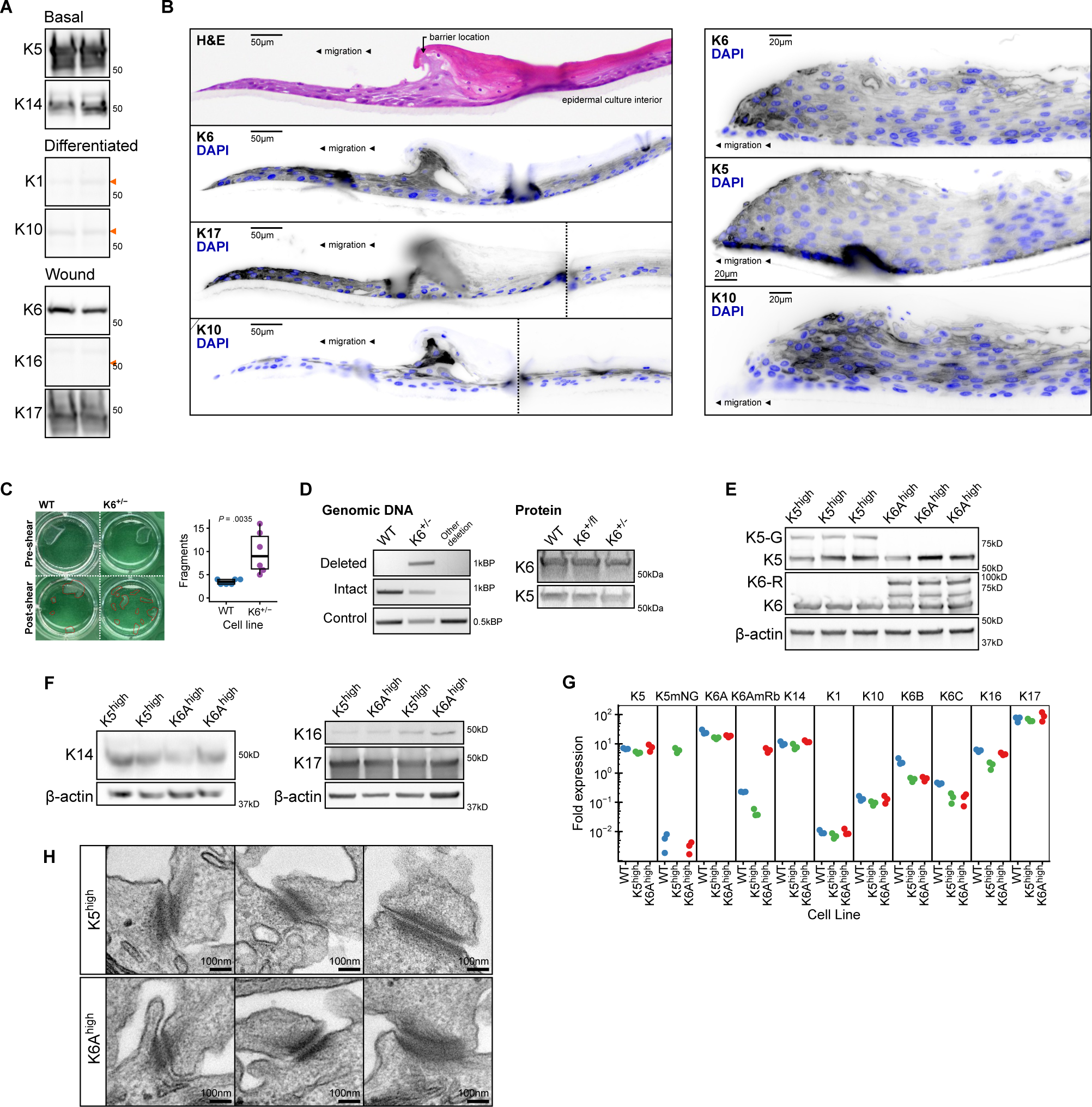
Keratin expression in keratinocyte culture models. A. Western blot analysis of keratin expression in human keratinocytes grown in submerged culture. Orange arrowheads indicate expected size for protein bands not clearly visualized. Samples from two independent cultures are shown. B. Additional examples of the epidermal culture wound model. Hematoxylin and eosin (H&E) staining or immunofluorescence labeling of wound-associated K6 and K17, steady-state basal K5, or steady-state differentiated K10. Left, migration edge; Right, epidermal culture interior. Dashed vertical lines, image re-alignment correcting for folds in the tissue section. See also Figure 1I. C. Mechanical stability of keratinocyte monolayers grown from wild-type or K6 heterozygous deletion (K6^+/-^) keratinocyte cell lines measured using a dispase-based fragmentation assay (see Methods). *n* = 6 monolayers per group. D. Genomic DNA PCR and Western blot analysis confirming heterozygous deletion of the type-II wound-associated keratin cluster (KRT6A/B/C), resulting in partial reduction of K6 protein (see Methods). K6^+/-^, heterozygous deletion clone; K6^+/fl^, keratinocytes with floxed KRT6A/B/C genomic cluster, prior to Cre-mediated recombination. E. Western blot analysis of endogenous (K5, K6) and exogenously expressed (K5-G, K6-R) keratins in K5^high^ and K6A^high^ cell lines (see also Figure 2B). F. Western blot analysis of other endogenously expressed keratins in K5^high^ and K6A^high^ keratinocyte cell lines. G. Keratin transcript levels in parental (WT), K5^high^, and K6A^high^ keratinocyte cell lines measured by qRT-PCR. Individual keratin transcript levels were compared to the average level of all keratin transcripts measured. *n* = 3 samples per group. H. Thin-section transmission electron microscopy images of typical desmosomes with keratin intermediate filament insertions in K5^high^ and K6A^high^ keratinocyte cell lines.

**Figure S2.**
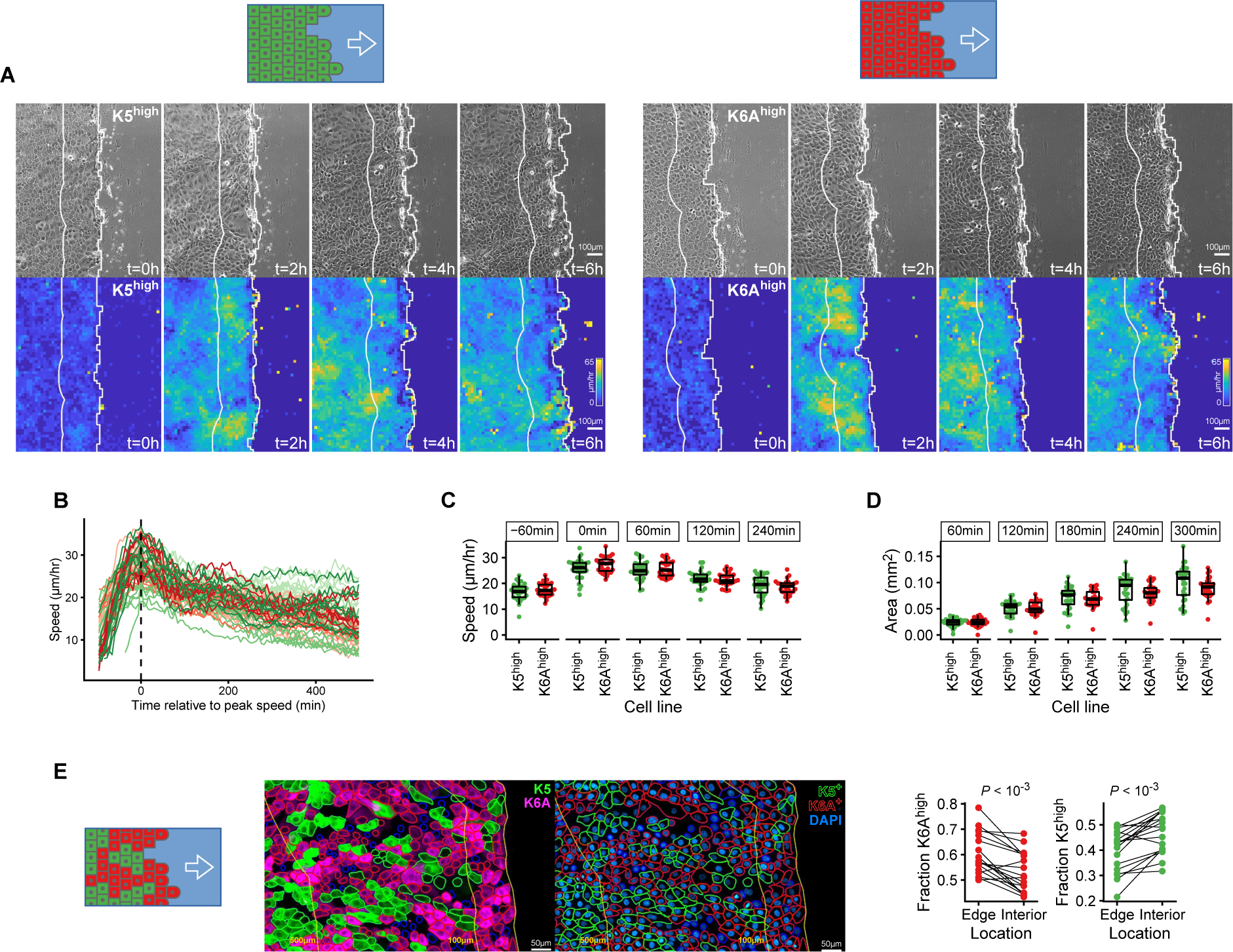
Wound-associated K6A supports keratinocyte migration in monolayers. A. Uniform-population monolayers of K5^high^ or K6A^high^ cells were scratched with a pipette tip to create a wound and imaged live as cells migrated into the wound area. The migration front was tracked using a computer vision pipeline and local and global migration speeds were calculated (see Methods). Top, phase-contrast images. Bottom, local migration speed maps. White lines, 200-μm wide band tracking the wound edge. See also Video 2. B. Time courses of average local migration speed in the area 10-μm to 200-μm from the wound edge for individual scratch wounds were aligned by the time of peak migration speed. Green, K5^high^ cells; red, K6A^high^ cells. *n* = 28-30 scratch wounds per group. C-D. Comparison of average local migration speed (C) and area closed (D) for migrating K5^high^ and K6A^high^ monolayers over time. In (C), time windows are aligned relative to the time of overall peak migration speed for each monolayer. *n* = 28-30 scratch wounds per group. E. Mixed population monolayers containing both K5^high^ and K6A^high^ cells were scratched to create a wound, then fixed and imaged after 24-hours of migration. Cells were segmented and classified using a computer vision pipeline (center; see Methods), then the proportion of K5^high^ and K6A^high^ cells in the 100-μm band closest to the wound edge was compared to the next 400-μm. *n* = 18 scratch wounds.

**Figure S3.**
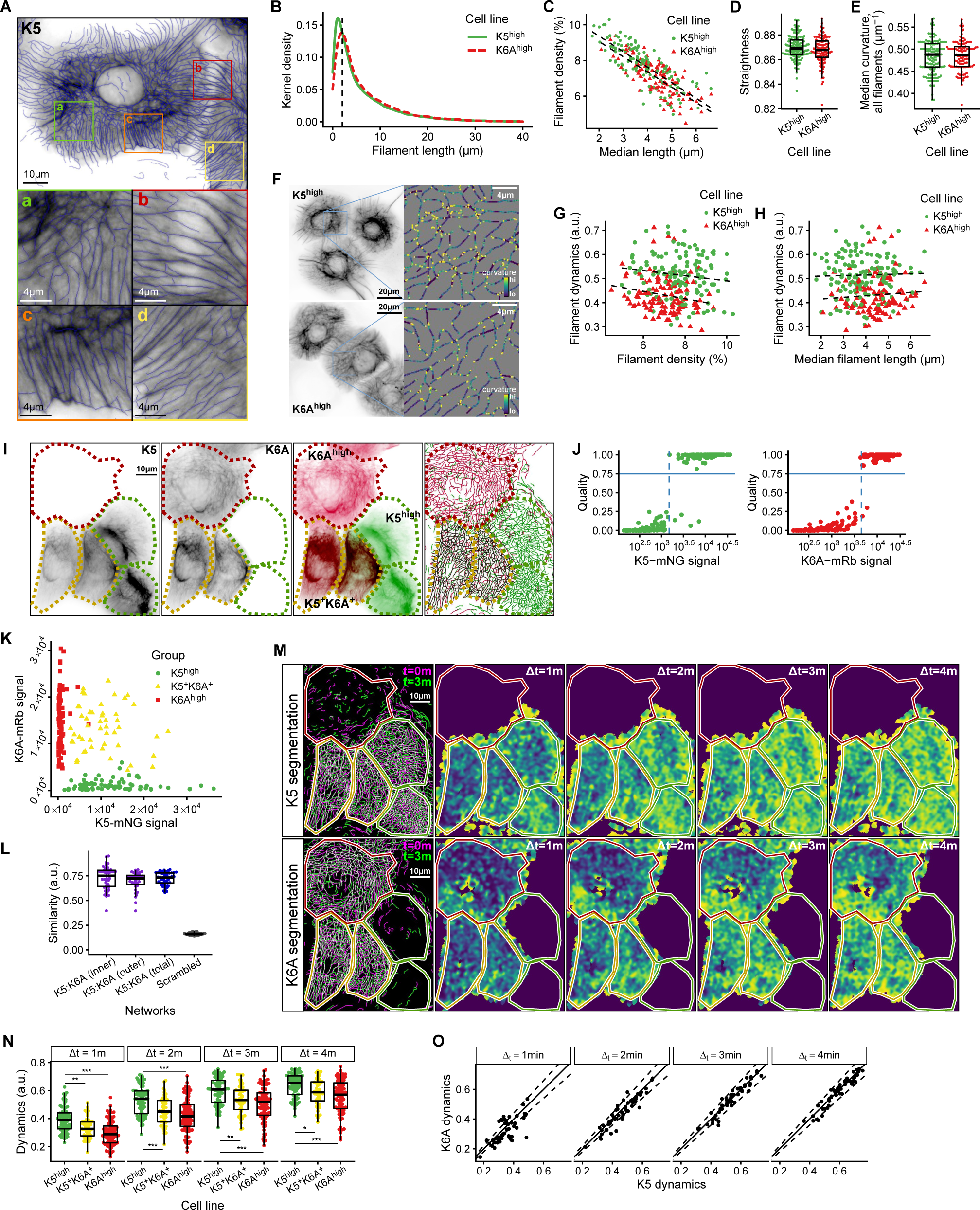
Wound-associated K6A alters filament network dynamics, but not static architecture. A. Demonstration of the filament segmentation pipeline (see Methods). Network segmentation (blue lines) overlayed on fluorescence imaging of K5-mNeonGreen. B-H. Images of keratin filament networks in K5^high^ and K6A^high^ cells were segmented and compared. This dataset comprises the first frame of each image sequence analyzed in Figure 5A-E. *n* = 110-139 cells per group with 1.7×10^4^-2.6×10^4^ total filaments per group. B. Length distribution of individual segmented filament lengths in K5^high^ and K6A^high^ cells. C. Correlation between filament density, defined as the percentage of cell area pixels included in the segmented filament network, and median filament length per cell. Dotted lines, linear regression models (K5^high^, *R^2^_adj_* = .61; K6A^high^, *R^2^_adj_* = .56). D. Median filament straightness per cell, defined as the ratio between the end-to-end distance and length of a filament. E. Median filament network curvature, defined as the median of point curvature estimates at each pixel included in the segmented filament network. This metric limits the influence of short filaments on the summary statistic. F. Example images of filament point curvature estimates. G. Correlation between filament network dynamics (3-minute time interval) and filament density. Dotted lines, linear regression models (K5^high^, *R^2^_adj_* = .01; K6A^high^, *R^2^_adj_* = .03). H. Correlation between filament network dynamics (3-minute time interval) and median filament length. Dotted lines, linear regression models (K5^high^, *R^2^_adj_* = <0.01; K6A^high^, *R^2^_adj_* = <0.01). I-O. Analysis of keratin filament network dynamics in a mixed culture of K5^high^, K6A^high^, and K5^high^/K6A^high^ (K5^+^K6A^+^) cells. *n* = 49-87 cells per group in 68 total movies. I. Two-channel fluorescence imaging of tagged K5 and K6A in the mixed cell population. The filament networks were separately segmented on each channel (right). Dotted outlines indicate cell classification: green, K5^high^; red, K6A^high^; yellow, K5^high^/K6A^high^ (K5^+^K6A^+^). J. Method for cellular keratin level determination. For each channel, cell filament segmentation quality scores, defined as the fraction of cell area with a dynamics score match at a 1-minute interval (see Methods), compared to the median fluorescence signal. A high score is synonymous with high fluorescent tag expression. A quality score of 0.75 (solid horizontal lines) was used to separate high from low expression levels of the keratin associated with each channel. The quality score showed better separation power than cut-offs defined by median fluorescence signal (dashed vertical lines). K. Median K5 and K6A fluorescence signals in cells classified as K5^high^, K6A^high^, and K5^high^/K6A^high^ (K5^+^K6A^+^) based on the segmentation quality score. L. For K5^high^/K6A^high^ cells (*n* = 49), the filament networks segmented separately based on the K5 and K6A signals were compared for similarity, defined as 1 minus the dynamics score, to validate the segmentation pipeline. To confirm even mixing of each isoform throughout the cell, similarity scores (K5:K6A) were averaged across the outer half (closest to the membrane), inner half, and total area of each cell. Synthetic filament networks created by applying small random shifts and rotations to the actual segmented networks while preserving local network densities (Scrambled) were used as a control (see Methods). M. Filament segmentation and network dynamics score maps for K5^high^, K6A^high^, and K5^high^/K6A^high^ cells. Outlines indicate keratin expression levels as in (I). N. Comparison of network dynamics scores between cell types. *, *P* < .05; **, *P* < .01; ***, *P* < 10^-3^. O. Correlation between network dynamics scores calculated separately for the K5 and K6A channels in K5^high^/K6A^high^ cells. The solid diagonal line indicates equal scores, the expected result for evenly intermixed keratins in a single network with perfect segmentation. Dashed lines indicate ±10% deviation. *n* = 49 cells in 28 movies.

**Figure S4.**
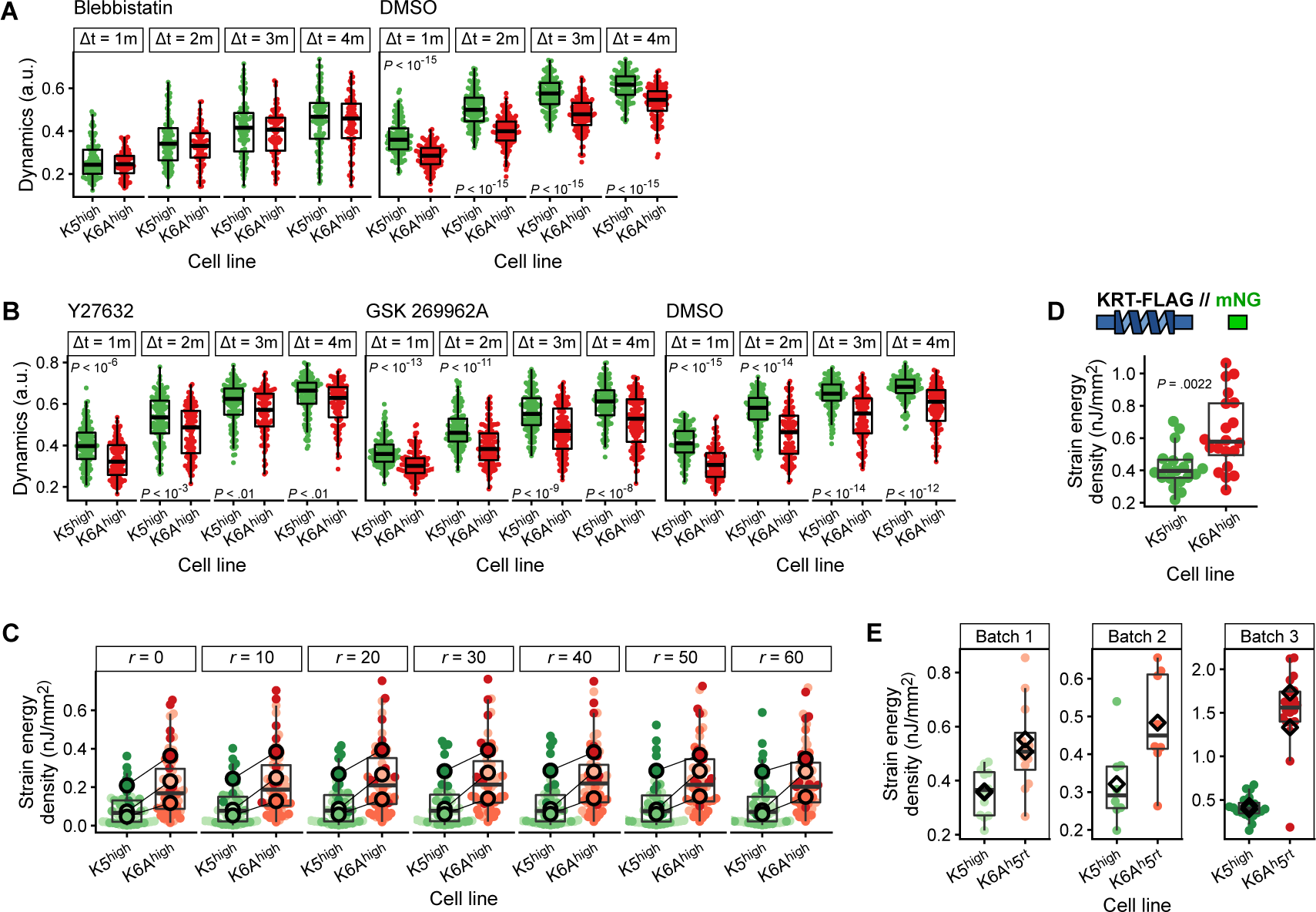
Network dynamics under reduced myosin motor activity and sensitivity analysis of traction force microscopy experiments. A-B. Keratinocytes were treated with 30 μM of the myosin ATPase-inhibitor Blebbistatin or DMSO control for 1 hour (A), or with 5 μM or 1 μM of the ROCK inhibitors Y27632 and GSK 269962A respectively, or DMSO control, for 2 hours (B). After drug incubation, keratin network dynamics were measured and compared between cell lines. A, *n* = 81-143 cells in 27-45 movies per group. B, *n* = 110-144 cells in 18-20 movies per group. C. Strain energy density of K5^high^ and K6A^high^ cells measured using traction force microscopy with different expansion radii of the cell area mask beyond visible tagged-keratin fluorescence signals. Expansion radius in pixels. See also Figure 6A. *n* = 51-58 cells per group. D. Strain energy density of K5^high^ and K6A^high^ cells from alternate cell lines without fluorescent tags linked to the exogenously expressed keratins. *n* = 19-20 cells per group. E. Strain energy density of cells expressing a chimeric keratin containing the head domain of K6A joined to the rod and tail of K5 (K6A^h^5^rt^) or full-length K5, with results separated by TFM substrate batches. Diamonds indicate mean strain energy density for cells on individual TFM substates (1-2 per group per batch). See also Figure 6B. *n* = 40 cells per group.

**Figure S5.**
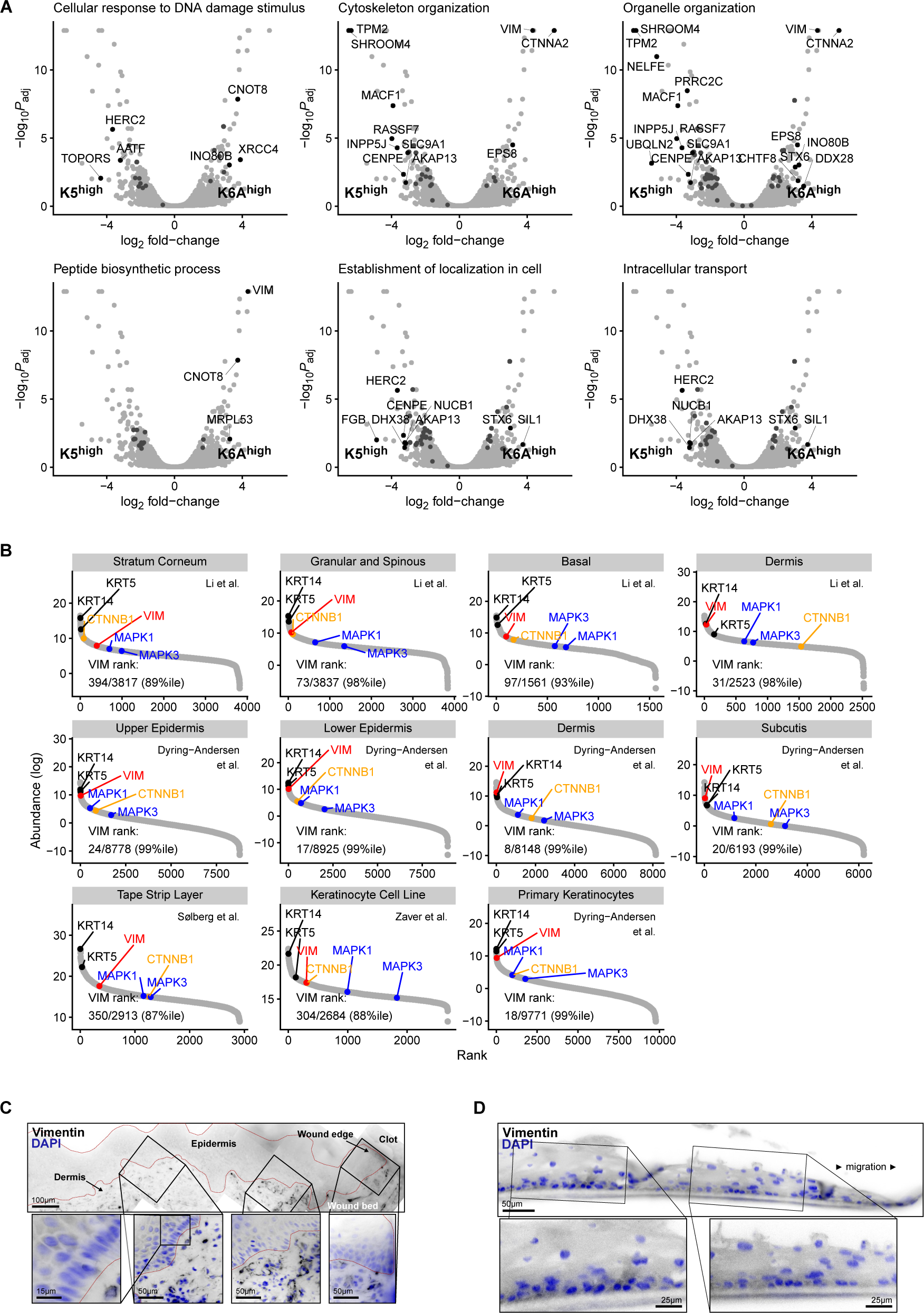
Other proteins differentially associated with keratin isoforms and vimentin expression in keratinocytes. A. Volcano plots highlighting differentially expressed proteins within the most enriched Gene Ontology Biologic Process labels. See also Figure 6C. B. Abundance and rank of selected proteins detected in published proteomics datasets of human skin samples, primary keratinocytes, and a keratinocyte cell line. Vimentin (VIM) is highlighted in red. Selected keratins (KRT5 and KRT14) are highlighted in black. β-catenin (CTNNB1), which has both a structural role in epithelia as a key component of adherens junctions as well as a signaling role, is highlighted in orange. Signaling molecules ERK1 (MAPK3) and ERK2 (MAPK1) are highlighted in blue. C. Immunofluorescence labeling of vimentin in a human skin wound. The wound edge is at the right. The epidermis is outlined in red. Note indistinct labeling of keratinocytes in the epidermis compared to strong labeling of fibroblasts and endothelial cells in the underlying dermis. D. Immunofluorescence labeling of vimentin in the epidermal culture wound model. The migration edge is at the right.

**Figure S6.**
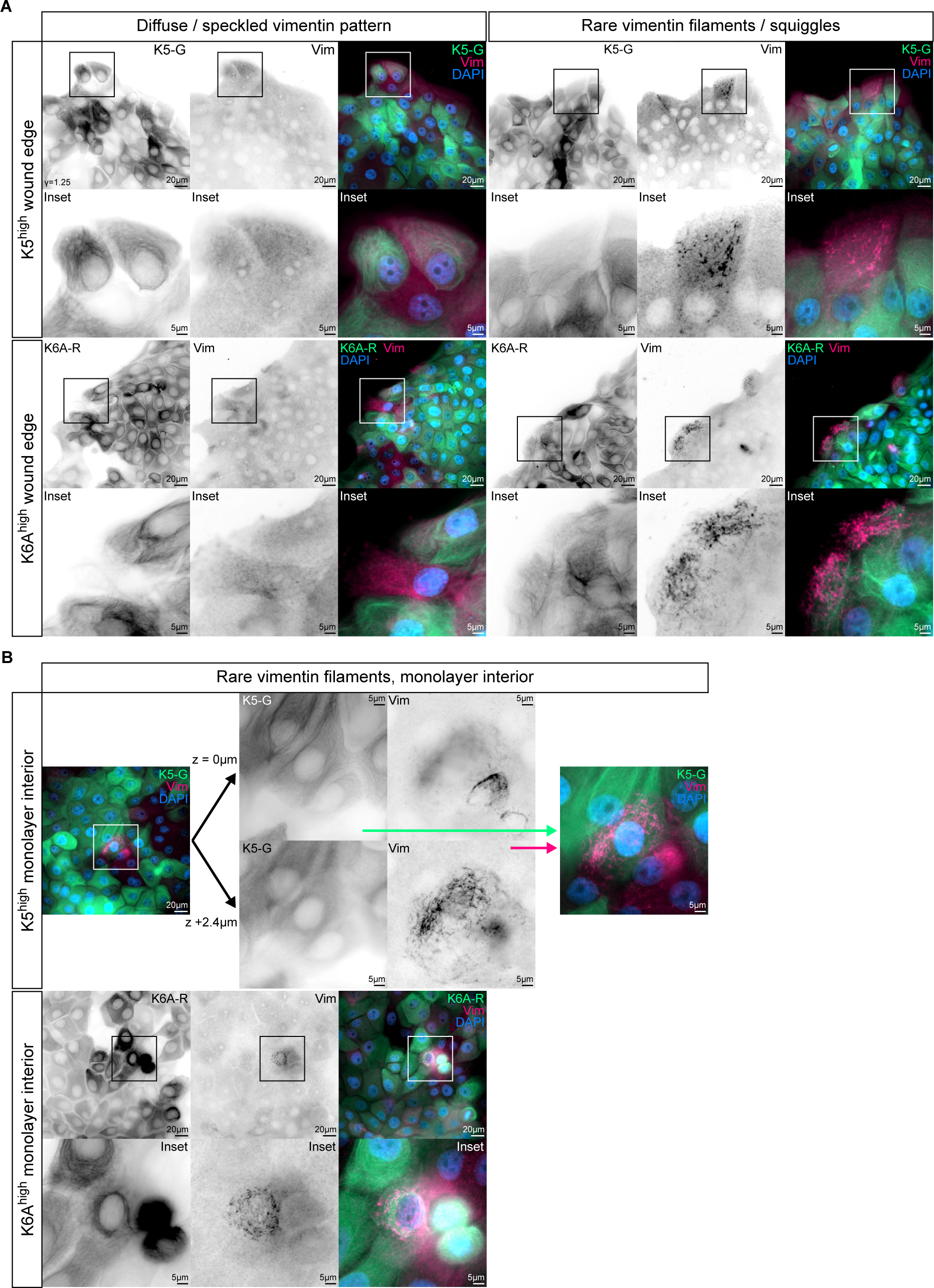
Non-filamentous vimentin is present throughout keratinocyte monolayers. A. Immunofluorescence imaging of vimentin (Vim) in monolayers of K5^high^ and K6A^high^ cells. Keratins are detected by their associated fluorescent tags (K5-G, K5-mNeonGreen; K6A-R, K6A-mRuby2). Cell monolayers were scratched to create a wound area 24-hours prior to fixation and processing for imaging. Diffuse or speckled vimentin staining patterns (left columns) were typical. Cells displaying vimentin squiggles or filaments were extremely rare (right columns, not quantified). B. Additional images highlighting rare cells away from the scratch wound edge with detectable vimentin squiggles or filaments. Top row center insets compare keratin and vimentin images in different focal planes. Top row right inset overlays keratin and vimentin images from different focal planes with the sharpest focus.

**Figure S7.**
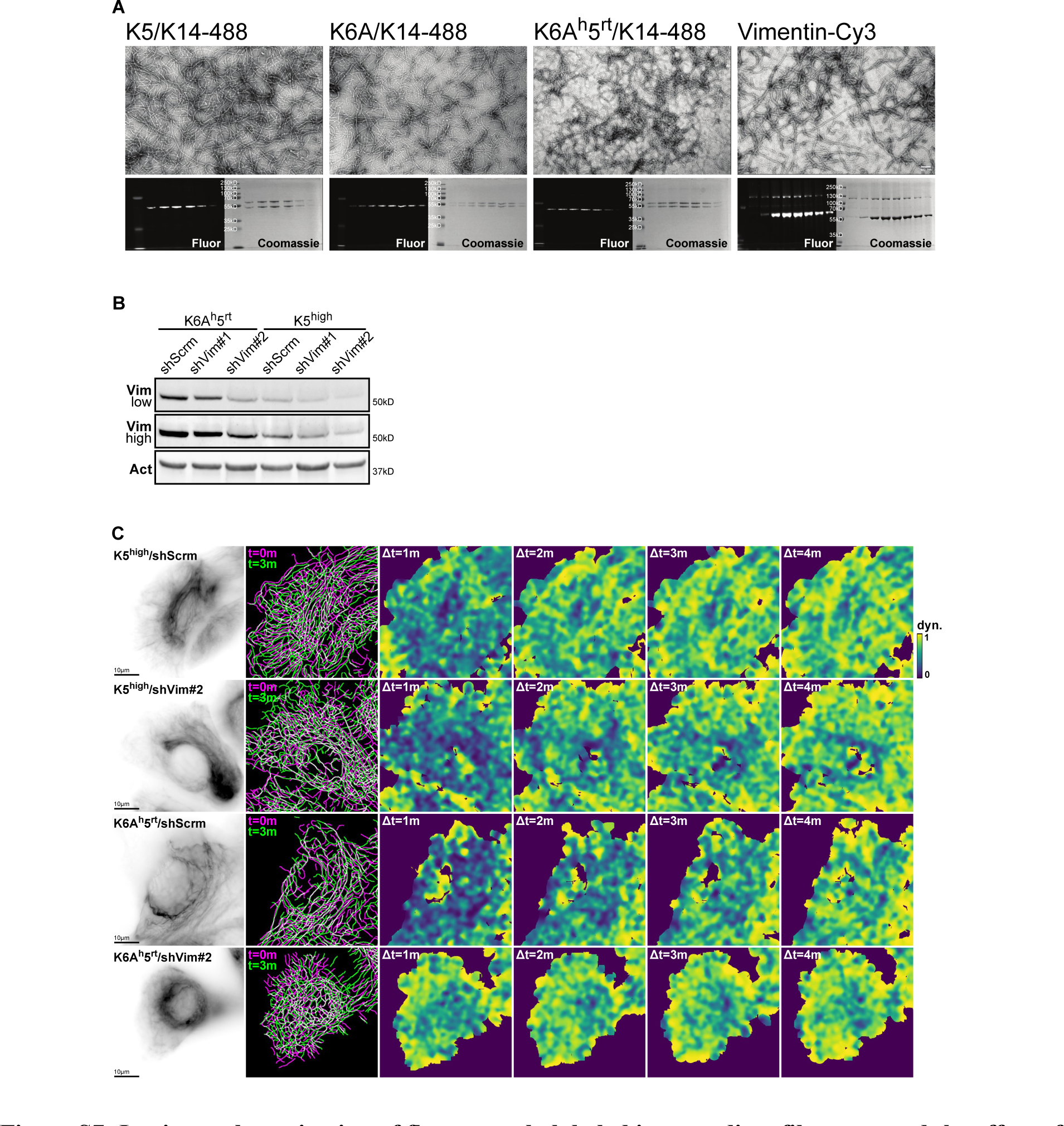
In vitro polymerization of fluorescently labeled intermediate filaments and the effect of vimentin knockdown on keratin filament dynamics. A. Negative stain electron microscopy and gel electrophoresis analysis of in vitro polymerized fluorescently labeled K14 (K14-488) with K5, K6A, or K6A^h^5^rt^ chimera; and in vitro polymerized fluorescently labeled vimentin (Vimentin-Cy3), confirming that fluorescent labels do not interfere with polymerization. For keratins, the Coomassie-stained gel images show the heterotypic complexes purified in urea, and the corresponding fluorescence images show K14-488. For vimentin, the Coomassie-stained gel image and the corresponding fluorescence image show that vimentin-Cy3 is able to form homodimers in the presence of less than 2 mM guanidinium chloride. See Methods. B. Western blot analysis of vimentin knockdown by shRNA (shVim#1, shVim#2 versus scrambled control, shScrm) in keratinocytes expressing a chimeric keratin containing the head domain of K6A joined to the rod and tail of K5 (K6A^h^5^rt^) or full-length K5 (K5^high^). The vimentin blot is displayed twice, with high and low brightness, to account for the different baseline vimentin levels in K5^high^ and K6A^h^5^rt^ cells. See also Figure 7B for an alternate display of this blot. C. Keratin network dynamics scores in K6A^h^5^rt^ cells and K5^high^ cells expressing shRNA targeting vimentin (shVim#2) versus scrambled control (shScrm). Example keratin filament images, filament segmentations, and filament dynamics score maps. See also quantification in Figure 7I.

**Video 1. Wound-associated K6A supports migration in three dimensional epidermal cultures.**

Paired epidermal cultures of opposite cell lines initially separated by a 500-μm gap and imaged live over 24.6 hours at a sampling rate of 12 minutes per frame. Playback at 12 frames per second. Green and red lines indicate the farthest extent of migration for each culture. See also Figure 2H-K.

**Video 2. Uniform-population monolayer migration assays do not detect differences between K5^high^ and K6A^high^ cells.**

Uniform-population monolayers of K5^high^ or K6A^high^ cells were scratched to create a wound and imaged live over 12.5 hours at a sampling rate of 6 minutes per frame. Playback at 30 frames per second. The migration front was segmented using a computer vision pipeline and local migration speeds computed in blocks of 15-μm side-length by particle image velocimetry (see Methods). See also Figure S2A-D.

**Video 3. Mosaic monolayer migration assays reveal subtle differences between K5^high^ and K6A^high^ cells during migration.**

A mixed population monolayer containing K5^high^, K6A^high^, K5^high^/K6A^high^, and K5^low^/K6A^low^ cells were scratched to create a wound and imaged over 12.5 hours at a sampling rate of 6 minutes per frame. Playback at 30 frames per second. See also Figure 3A-F. Keratin expression regions in the monolayer were segmented based on the fluorescence signal from the tagged keratin constructs (left). Local migration speeds computed in blocks of 15-μm side-length by particle image velocimetry (right; see Methods).

**Video 4. Wound-associated K6A supports a transient migration advantage in monolayers.**

Pairwise comparison of average local migration speed between keratin expression regions relative to the global migration peak. Each point represents one scratch wound. Points located along the diagonal line indicate equal local migration speeds in the two keratin expression regions. Points above or below the diagonal indicate that one region is moving faster than the other. *n* = 36 scratch wounds. See also Figure 3E-F.

**Video 5. Wound-associated K6A supports a transient migration advantage in single cells.**

Individual keratinocytes were sparsely seeded and allowed to migrate without exogenous directional cues. Imaged over 18 hours at a sampling rate of 3 minutes per frame. Playback at 24 frames per second. See also Figure 3G-I.

**Video 6. Wound-associated K6A alters keratin filament dynamics.**

Keratin filaments in K5^high^, K6A^high^ (plays first), and K5^high^/K6A^high^ (plays second) cells were imaged live over 24 minutes at a sampling rate of 30 seconds per frame. Playback at 24 frames per second. Filament networks were segmented using a computer vision pipeline (see Methods). Filament segmentations are overlayed on the tagged keratin fluorescence images. See also Figures 4A-E and S3I-O.

